# Two melanic pigment patterns are associated with a sex chromosome-linked oncogene in the mountain swordtail *Xiphophorus nezahualcoyotl*

**DOI:** 10.64898/2026.06.08.730778

**Authors:** Lyle A. Given, Landen Gozashti, John J. Baczenas, Rhea Sood, Nadia B. Haghani, Kelsie E. Hunnicutt, Theresa R. Gunn, Gabriel A. Preising, Peter H. Sudmant, Daniel L. Powell, Tristram O. Dodge, Molly Schumer

## Abstract

Sex-linked traits are widespread, but their genetic architecture has been challenging to characterize, due in part to the repetitive and structurally complex nature of sex chromosomes. In swordtails and platyfish of the genus *Xiphophorus*, diverse melanic pigmentation patterns are thought to be controlled by a region on the sex chromosomes classically referred to as the “macromelanophore determining locus”. Despite nearly a century of study, the identity of the causal gene remains controversial, partially due to previous inability to fully sequence the sex chromosomes. Here, we characterize and investigate two melanin-based pigmentation phenotypes in the species *X. nezahualcoyotl*: “spotted side” and “marmoratus”. We generate a gapless near-telomere-to-telomere *X. nezahualcoyotl* assembly and perform GWAS to identify regions associated with pigmentation pattern variation and sex-determination. We find both patterns map near the sex-determining region and to a narrow interval near the oncogene *xmrk*. By generating additional long-read assemblies of sex chromosomes derived from individuals with distinct phenotypes, we find haplotypes containing *xmrk* can be both X- and Y-linked, and vary dramatically in gene content, structure, and accumulation of repetitive elements including a newly described composite satellite. This variability may impact the region’s stability and affect recombination between haplotypes associated with each pattern. Our results shed light on a longstanding debate surrounding the genetic architecture of sex-linked phenotypes. More generally, we showcase how long-read sequencing can reveal phenotypic variation linked to complex and dynamic genomic regions, which may contribute to the evolution of diverse sex-linked traits.

## Introduction

Sex-linked traits have captured special attention in the field of evolutionary genetics due to a puzzling observation: males and females of a species can dramatically differ in phenotype despite sharing much of their genome. While these differences commonly manifest as sexual dimorphism [1,2], sex-linked traits can also occur as polymorphisms that segregate at different frequencies within each sex [3]. These two kinds of sex-linked traits can reflect distinct evolutionary processes, but a key driver in their evolution is likely sexual conflict, which arises when different phenotypes optimize fitness in males and females [4]. Theory predicts that conflict can be resolved through sex-specific gene expression (e.g. modulated by sex hormones) or through physical localization of the underlying genes to sex chromosomes [5,6]. Such restriction of sexually antagonistic genes to the sex chromosomes may also underpin cases of sex-limited polymorphisms maintained by balancing selection [3]. The genetic architecture of sex-linked traits is an active area of research, and phenotypes governed by these mechanisms have been shown to be diverse, ranging from sperm function [7] to coloration ornaments [8].

Studying the genetic architecture of sex-linked traits is complicated by the wide diversity of sex chromosomes, which can range from single-base pair variants to highly heteromorphic chromosomes [9]. Some of the best studied systems, such as the Y and W chromosomes of mammals and birds, are small and degenerate. This is because these sex chromosomes are ancient, and millions of generations of suppressed recombination and weaker efficacy of selection have allowed for the accumulation of repetitive sequences, including interspersed mobile element insertions and tandemly repeated satellite DNA sequences [10]. However, there is a wide spectrum of sex chromosome age and divergence across the tree of life [11]. Notably, Y chromosomes in some lineages are larger than their homologous X [12], resulting from gain of genes and repetitive DNA [10]. While researchers have made considerable progress in understanding the evolution of sex-linked traits in classic heteromorphic XY/ZW systems, less is known about how these evolve in species with less differentiated sex chromosomes. These systems, which are often younger and actively experiencing gene movement and repetitive element accumulation on the sex chromosomes, can provide a unique window into the interplay between traits and chromosome evolution [9].

A major technical hurdle in the study of the genomic architecture of sex-linked phenotypes is the sequence complexity and structural variation of sex chromosomes. Although next-generation sequencing is now widespread across many non-model species, assemblies based on short-read sequencing approaches have limited resolution in complex regions. Recent advances in long-read and ultra-long-read sequencing technologies and assembly algorithms now allow fully complete genomes to be generated for the first time [13–16]. However, due to the higher cost of long-read sequencing, a common approach remains to map short-read whole-genome resequencing data to a single reference genome. This approach has several limitations: sex chromosomes can be incompletely assembled or missing in reference genomes (especially if the heterogametic sex was chosen [17]), the data may be especially prone to mis-mapping and reference bias [13], and relying on a single reference cannot fully capture the variation present in these complex and highly dynamic regions [18]. Thus, the contribution of sex chromosomes to phenotypic variation has often been overlooked [3,19]. In humans, model organisms, and domesticated species, there has been an increasing application of long-read sequencing to represent the full spectrum of genomic variation within species [20,21]. However, in non-model taxa, it was cost-prohibitive to use long-read approaches to sequence and compare multiple individuals from the same population until recently [22], which has hampered progress in this area. Species with high rates of sex-linked polymorphisms and compact genomes represent powerful systems where long-read sequencing can be more broadly applied to reveal the contributions of these complex and dynamic regions to trait variation.

*Xiphophorus* is a phenotypically diverse genus of Poeciliid fishes native to eastern Mexico and Central America, whose genetics have attracted attention for nearly a century [23]. Species in this group display varying degrees of sexual dimorphism [24] and sex-limited polymorphism [25,26]. Coloration traits underpinned by diverse cell types (e.g. melanophores, iridophores, xanthophores, and erythrophores) are often sex-linked [27,28]. Based on cytogenetics and laboratory crosses, researchers have found that the sex chromosomes are generally homomorphic [29] and that the location of the sex determining region is largely homologous between species [30], with some notable exceptions [31,32]. However, it remains puzzling why there are so many sex-linked traits in *Xiphophorus* and how they have evolved. A further complication to understanding the genetic architecture and evolutionary history of sex-linked traits in *Xiphophorus* is that only the sex chromosomes of one species (*X. maculatus)* have been studied at the genomic level [33]. Importantly, the gene content, repetitive landscape, and structural variation between the sex chromosomes, as well as the identity and the physical location of the sex-determining locus, remain unknown in most species.

The genetic basis of sex-linked melanic patterns is a longstanding question in *Xiphophorus* genetics [23,34,35]. Across the genus, researchers have documented myriad patterns driven by large, melanin-producing cells called macromelanophores, which segregate in at least ten species [36]. Controlled crosses suggest that distinct macromelanophore patterns within and across species are usually driven by a series of codominant alleles at a sex-linked macromelanophore determining locus classically called Mdl [30,37, but see 38,39]. Notably, misregulation of macromelanophore patterns can lead to malignant melanomas in *Xiphophorus* hybrids [23,40], and melanomas also occasionally occur in pure species [41–43]. Melanoma formation in hybrids results from misregulation of the gene *xmrk* (*<u>X</u>iphophorus <u>M</u>elanoma <u>R</u>eceptor Tyrosine-<u>K</u>inase*) [44–46], a polymorphic sex-linked oncogene that originated 5-10 million years ago via duplication of the epidermal growth factor receptor gene *egfrb* [47]. After its discovery, *xmrk* was proposed to also drive macromelanophore presence in *X. maculatus* (that is, *xmrk* is Mdl [44]). However, later work by Weis & Schartl [48] proposed an alternative model whereby *xmrk* is instead tightly linked to Mdl and enhances existing macromelanophore patterns, but *xmrk* presence does not drive presence of macromelanophores. Key evidence for this assertion was that two *Xiphophorus* species, *X. hellerii* and *X. nezahualcoyotl*, have macromelanophore patterns but lacked *xmrk* based on Southern Blot analysis [48]. However, later work cast some doubt on this assertion. Fernandez and Bowser [43] documented malignant melanomas in non-hybrid *X. nezahualcoyotl*, leading to speculation that this species may be polymorphic for *xmrk*, and analysis of short-read Illumina data from a single *X. nezahualcoyotl* sample found evidence of *xmrk* [49]. However, the genetic architecture of macromelanophore patterns in *X. nezahualcoyotl* and their potential associations with *xmrk* have yet to be resolved.

Here, we address these longstanding questions by investigating the genetic architecture of sex-linked macromelanophore patterns in *X. nezahualcoyotl*. We characterize two macromelanophore phenotypes, “spotted side” and “marmoratus,” the latter of which was previously undescribed in this species. We use GWAS to identify narrow *xmrk*-containing haplotypes associated with each trait, which are adjacent to the sex-determining region. By comparing contiguous sex chromosome assemblies from individuals with diverse phenotypes, we further investigate coding and non-coding variation that may contribute to macromelanophore pattern variation and chromosome evolution. Our findings reveal that these traits are associated with a complex and dynamic genomic region, highlighting the challenges, but also new possibilities, of linking genes on the sex chromosomes with phenotypic variation.

## Results

### Two spotting phenotypes segregate in X. nezahualcoyotl from the Río Tanchachín

The mountain swordtail (*X. nezahualcoyotl*) is a member of the Northern Swordtail clade, endemic to the Río Pánuco and Río Tamesí drainages in Eastern Mexico [50]. Prior work has primarily studied populations from the Río El Salto, part of the Pánuco drainage [43,48, 51] and the Río Gallitos in the Tamesí drainage [49,52]. Here, we collected *X. nezahualcoyotl* from a natural population in the Río Tanchachín, in the southern extent of its range (Fig. 1a). While sampling fish from the river, we observed the apparent segregation of two distinct patterns (Fig. 1b). One pattern manifested as black spots distributed across the flank, ranging from 4-6 distinct rows to an irregular blotchy arrangement, matching the description of a previously described sex-linked pattern known as “spotted side” [51]. The other pattern was made up of larger, irregular blotches restricted to the ventral flank. This pattern resembled “marmoratus,” a trait that had been previously described in a related species *X. montezumae* [48]. For consistency with previous work, we refer to this marmoratus-like pattern in *X. nezahualcoyotl* as “marmoratus” due to its visual similarity, but we note that we are unable to determine its homology since the genetic basis of marmoratus in *X. montezumae* has not been determined.

**Figure 1.**
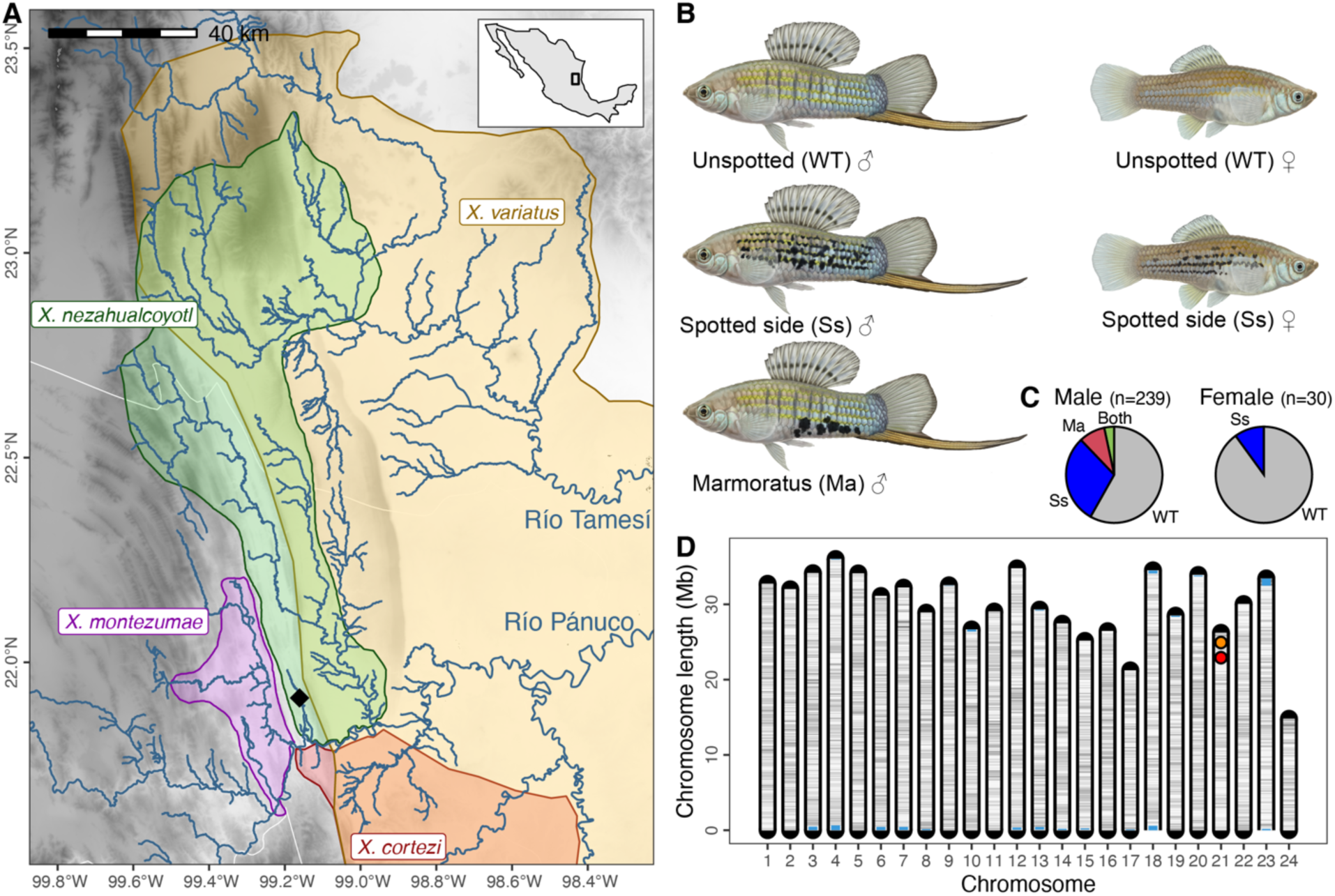
*X. nezahualcoyotl* from the Río Tanchachín segregate for multiple spotting patterns and a putative *xmrk.* **(a)** *Xiphophorus* species ranges in the Río Pánuco and Río Tamesí drainages within the Sierra Madre Oriental in Eastern Mexico. Map background is colored by elevation (white-low, black-high), with main rivers shown in blue. The mountain swordtail (*X. nezahualcoyotl*) is sympatric with the variable platyfish (*X. variatus*) in the northern extent of its range and borders the Montezuma swordtail (*X. montezumae*) and delicate swordtail (*X. cortezi*) in the south. Samples for this study were collected at the Río Tanchachín (black diamond) in the southern extent of the range. **(b)** In the Río Tanchachín, two classes of polymorphic spotting patterns segregate in males (n=239) and females (n=30), matching descriptions of the patterns “spotted side” and “marmoratus.” Spotted side and marmoratus can co-occur in males, but marmoratus appears to be absent in females. Illustrations by Dorian Noel. **(c)** Phenotypic frequencies estimated in a random sample of fish from the Río Tanchachín. Note that we have never observed marmoratus in female *X. nezahualcoyotl*. **(d)** A gapless, near complete genome from a female with spotted side. Chromosomes are shaded by density of coding sequence in 50 kb windows (white-low, black-high). Caps on the end of chromosomes represent telomeres. 22 of 24 expected chromosomes are telomere-to-telomere (T2T) while chromosomes 18 and 23 end instead in an rDNA array (shown in blue). The primary assembly of chr-21 contains two copies of an *xmrk*-like gene, representing a putative *xmrk* (shown in red; position 22.9 Mb) and its paralog *egfrb* (shown in orange; position 24.9 Mb).

We estimated phenotypic frequencies of spotted side and marmoratus in a subset of adults that were sampled randomly with respect to phenotype. Males (n=239) and females (n=30) were polymorphic for spotted side at frequencies of 33% and 10%, respectively. We observed marmoratus in males at 12% frequency and did not observe it in females (Fig. 1c). We note that, despite the small number of females included in this analysis, we have sampled hundreds of females at this site over three years and have never observed marmoratus in this sex. In males, we occasionally observed co-occurrence of spotting patterns within the same individual (Fig. 1c, Fig. S1).

### A gapless chromosome-level X. nezahualcoyotl assembly contains an xmrk-like gene

Due to a lack of genomic resources in this species, we first aimed to generate a high-quality reference assembly. We sequenced a wild-caught *X. nezahualcoyotl* female (hereafter Ss-F01) with the spotted side phenotype using Oxford Nanopore Technologies (ONT), generating an ultra-long-read dataset (57.2 kb read N50) at ∼80× coverage (Table S1). Assembly with hifiasm [53] and orientation with RagTag [54] resulted in a 715 Mb gapless primary assembly with 22 telomere-to-telomere (T2T; Fig. 1d) contigs out of 24 expected chromosomes. The two remaining long contigs each ended in one telomere and in one rDNA array (Fig. 1d). Other quality metrics, including BUSCO [55] and Merqury QV [56], indicated the assembly was highly complete with low error rates (Table S2). To annotate this new reference genome, we lifted over NCBI features from the congeneric species *X. maculatus* with LiftOn [57], which has been intensively studied and thus has high-quality annotations. Predicted proteins in *X. nezahualcoyotl* had high BUSCO completeness values, comparable to NCBI annotations in *X. maculatus* (Table S3), suggesting good quality of our LiftOn approach. To annotate repetitive content, we generated a pan-TE library of polymorphic elements in *Xiphophorus* with Pantera [58] followed by manual curation. This approach, which leverages presence-absence polymorphisms to identify candidate TEs, yields higher-confidence and less fragmented models compared to fully homology-based methods [58], but it may not capture transposable elements that are no longer segregating in the species group (e.g., ancient elements that may be no longer active). 19.73% of the *X. nezahualcoyotl* reference genome was annotated as repetitive using this curated repeat library (Table S4).

With a gapless and near-complete reference genome, we first aimed to ask if the *xmrk* gene was present in the reference. An inspection of the unitig graph suggested that the ultra-long ONT reads had permitted complete phasing of most chromosomes (Fig. S2). We therefore searched for *xmrk* within our gene annotations in both hifiasm “pseudohaplotypes” of chromosome 21 and found one contained a putative *xmrk* and *egfrb* (22.9 Mb and 24.9 Mb; Fig. S2) while the other only had *egfrb* (25.4 Mb). We also conducted a BLASTn search [59] for *xmrk* and found two hits on hap1 and one on hap2, consistent with the results of our gene annotations and with the known high similarity of *xmrk* and *egfrb.* Thus, the results of the gene annotations and BLASTn search provided preliminary evidence that *xmrk* is present and polymorphic in *X. nezahualcoyotl*, which we later confirmed with additional analyses.

### Spotted side and marmoratus show distinct spatial organization and consist of macromelanophores

Since the two patterns we study appeared to fall into distinct phenotypic classes, we hypothesized the melanic spots would show distinct spatial localization along the fish’s flank. To quantify spatial variation, we generated binary pattern masks using a modified version of the recolorize-patternize workflow [60]. We found that spotted side occurs broadly across the body and is centered around the lateral line (Fig. 2a) with patterns varying from a more lineated arrangement to larger blotches (Fig S3). In contrast, the location of spots in marmoratus is concentrated on the ventral flank, located primarily between the pelvic fin and caudal peduncle (Fig. 2a). When we analyzed the pattern masks with principal component analysis (PCA; Fig. 2b), two distinct phenotypic clusters formed, separating along PC1 (15.877%). The cluster of individuals with the spotted side phenotype further separates along PC2 (6.537%). This separation persisted when the groups were downsampled to an equal number of individuals (Fig. S3).

**Figure 2.**
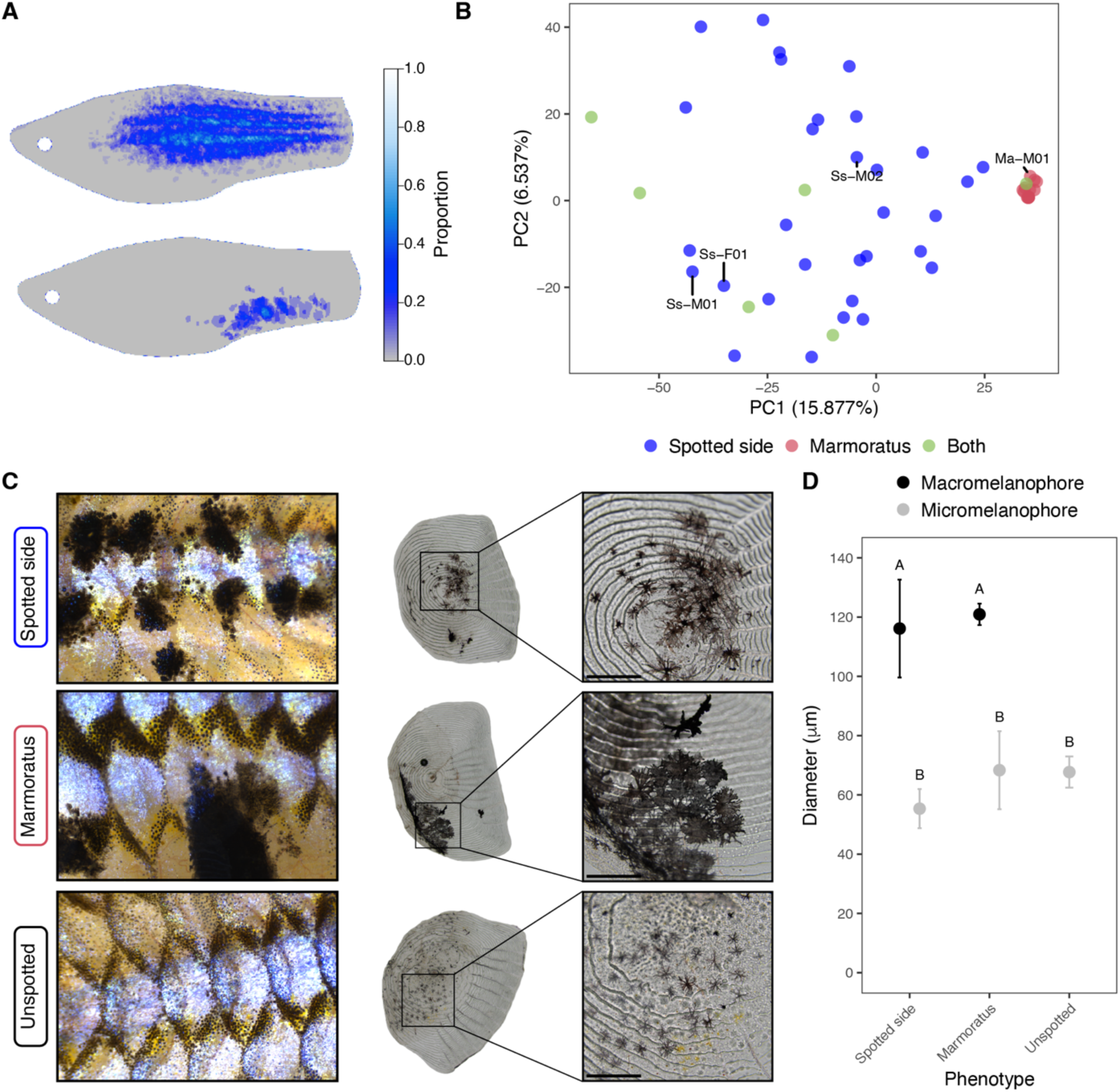
Macromelanophore patterns are similar at the cellular level but display spatial variation. **(a)** Heatmaps of individuals with spotted side (top) and marmoratus (bottom) generated from summed pattern raster objects. **(b)** A principal component analysis based on pattern raster objects shows two distinct clusters, corresponding to spotted side and marmoratus, that separate along PC1 (15.877%). Labels denote spotted individuals that were subsequently sequenced with long-reads. **(c)** Individuals with each phenotype were imaged at 2x magnification, and scales were imaged at 4x and 20x magnification. Scales from individuals with spotted side and marmoratus show macromelanophores, and the scale from an unspotted individual shows micromelanophores. The scalebars on the 20x magnification images are equal to 200 μm. Images were rotated when necessary and cropped to aid visualization. **(d)** Melanophore cell size results indicate macromelanophores are significantly larger than micromelanophores (two-way ANOVA, Tukey’s HSD, p=4e-7) but do not significantly differ between phenotypes (p=0.98). A and B labels denote statistically different groups based on Tukey’s HSD.

In *Xiphophorus*, two classes of melanic pigmentation cells—macromelanophores and micromelanophores—are distinguished by size, with cells greater than 100 µm considered macromelanophores [61]. Melanophores comprising both traits appeared localized in the dermis and epidermis, and we imaged scale-associated melanophores from spotted and unspotted regions of the body. We compared the diameter of spot-associated melanophores to the melanophores making up the reticulated “stipple” pigmentation associated with the edges of scales, which are expected to be micromelanophores (Fig. 2c). Scales plucked from regions with spotting patterns contained melanophores greater than 100 µm, confirming that both spotted side and marmoratus are underpinned by macromelanophores (Fig. 2d). We did not find evidence for a significant difference in cell diameter between macromelanophores from spotted side and marmoratus (Tukey’s HSD, p=0.98). Across spotted and unspotted individuals, we confirmed that the melanic cells not associated with these two spotting patterns were consistently smaller, ∼60 µm in diameter (i.e. micromelanophores). The macromelanophore cells comprising the two spotted patterns were consistently larger than the micromelanophore cells examined (Fig. 2d, two-way ANOVA, Tukey’s HSD, p=4×10^−7^).

### GWAS identifies a sex chromosome-linked association for macromelanophore patterns

Prior crossing experiments indicated that the spotted side macromelanophore pattern is sex-linked in *X. nezahualcoyotl* [51]; however, past work has not attempted to link genotype to spotting phenotype in this species and the role of *xmrk* remains unknown. Using low-coverage whole genome sequencing from a single population, we performed case-control GWAS to identify allele frequency differences between phenotypes. We used the near-T2T primary genome assembly of a female with spotted side as the reference genome for each pattern GWAS analysis. For the spotted side GWAS, our sample population consisted of 256 individuals with a phenotype frequency of 29%. Since we only observed marmoratus in males, we excluded females from the marmoratus GWAS, so the sample population consisted of 214 males with a phenotypic frequency of 10%. Both pattern GWASes implicated a single locus on chromosome 21 (Fig. 3a,3b) which surpassed the genome-wide significance threshold estimated by simulation for each GWAS by (p<2.6×10^−9^ and p<2.14×10^−9^). The interval spanning SNPs significantly associated with spotted side was 4.8 kb and overlapped a non-coding region that contained no genes, while the region associated with marmoratus spanned 70.1 kb and overlapped two genes: the previously noted *xmrk*-like gene (LOC102228937) and a gene with unknown function (LOC111611186) with homology to a CCHC-type zinc finger. The lead SNPs for the spotted side and marmoratus GWASes were 68.2 and 67.4 kb upstream of *xmrk*, respectively. In both cases, significantly associated alternate alleles were at high frequency in the spotted side and marmoratus groups while being at low frequency or absent in the control groups (Fig. 3a,b), which together is consistent with a dominant and highly penetrant genetic architecture of both spotted phenotypes. We found no evidence of population structure that associated with phenotypic variation based on a genomic PCA analysis (Fig. S4).

**Figure 3.**
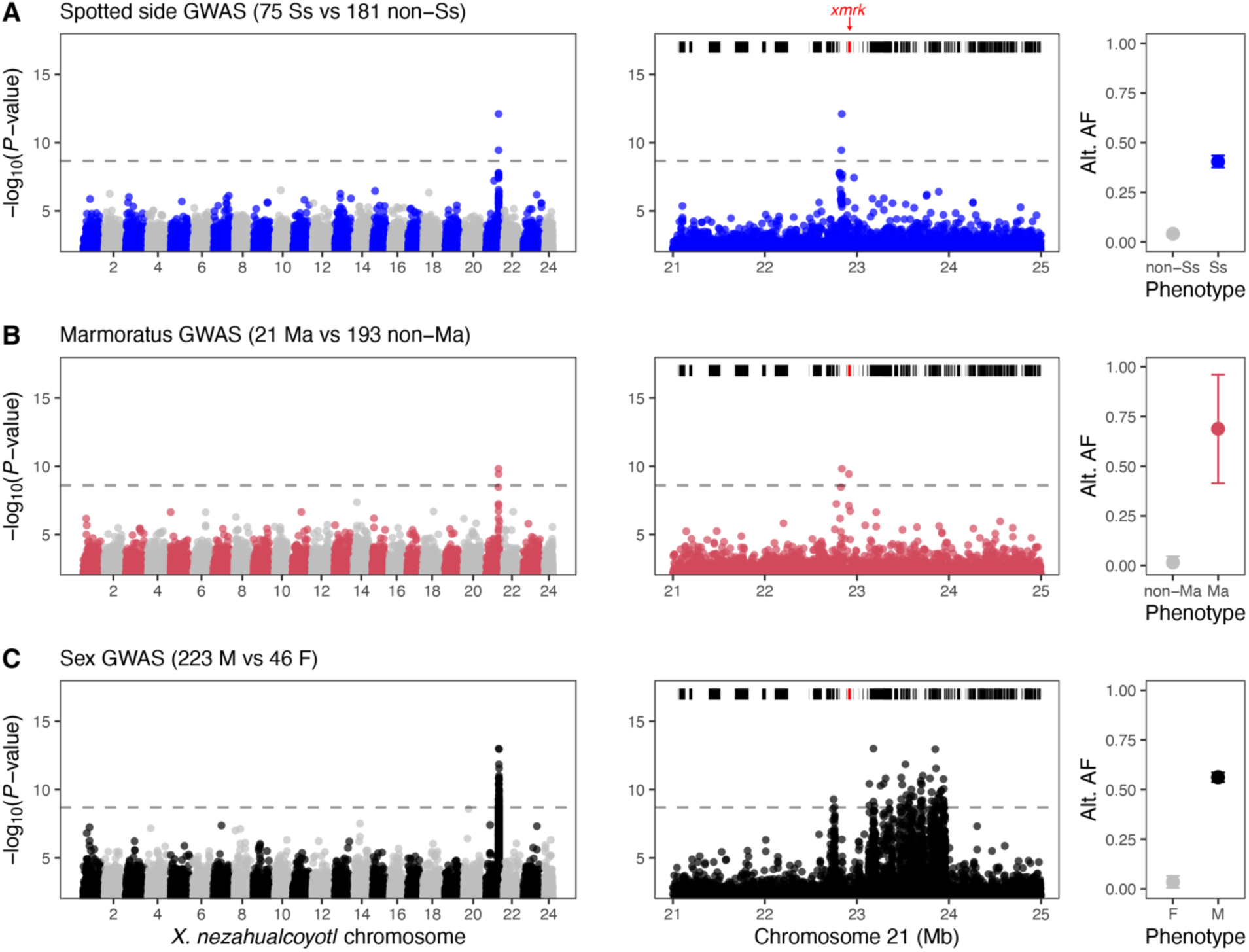
Case-control GWAS identify a region on chromosome 21 associated with spotted side **(a)**, marmoratus **(b)**, and sex **(c)**. Dashed lines show the 5% false positive threshold for each GWAS, based on 500 simulations. Left panel shows the whole genome, and middle panel shows a 4 Mb region surrounding the peak signal on chromosome 21 with annotations for protein-coding genes shown above (*xmrk* in red). Right panel shows the mean frequency of the alternative (non-reference) allele for significant SNPs within the GWAS interval from fish in different phenotype classes. Error bars denote ±2 standard error. Note that for the allele frequency analysis, we mapped to the alternate reference haplotype for chromosome (haplotype 2), which did not have *xmrk*, to aid in interpretation of the alternate allele frequency.

While chromosome 21 has been identified as the sex chromosome in *X. maculatus* [33], to our knowledge the sex chromosome in *X. nezahualcoyotl* has not been identified with genomic data. Accordingly, we performed a case-control GWAS using 223 males and 46 females to identify the sex-determining region. This GWAS identified a single locus on chromosome 21 surpassing the simulations-based genome-wide significance threshold (p<2×10^−9^). This region was 1.37 Mb wide and co-localized with the signal associated with the macromelanophore patterns (Fig. 3c). In this analysis, males had high frequency of alternate alleles at lead SNPs, consistent with a simple XY system with high penetrance. The GWAS results appear to suggest that ∼95% of the *X. nezahualcoyotl* sex chromosome recombines, and the variants underlying macromelanophore patterns localize near the sex determining region.

### Long-read sequencing of individuals with spotting patterns reveal association with the xmrk-like gene

Previous crossing results in *X. nezahualcoyotl* suggested the spotted side pattern is sex-chromosome linked [28,41]. However, across many *Xiphophorus* species, the sex-determining region is complex, structurally variable, and remains unassembled in most existing references [33,49]. To better capture patterns of sex-linked genetic variation associated with the spotting phenotype, we generated PacBio HiFi long-read assemblies from four additional males spanning a range of phenotypes: one unspotted male, one male with marmoratus, and two males with spotted side. We selected two spotted side males that displayed variability in their phenotype—one with a more lineated pattern and the other with a blotchier pattern (Fig. 2d; Fig. S5). The resulting hifiasm assemblies were highly contiguous and complete (Table S2). We also evaluated the quality of each chromosome 21 assembly using Merqury and found no evidence of misassembly near the sex-determining region (Table S5). We then annotated each assembly using the same workflow as described for the near-T2T female assembly.

Due to the high levels of structural complexity in the region as revealed by alignments (Fig. S6), we took an alternative approach to investigate other possible candidates for the macromelanophore patterns. To identify homologous sequences without relying on whole-genome alignments, where high levels of structural complexity made interpretation of alignments challenging, we took an alternative approach and performed GWAS to each long-read assembly “pseudohaplotype” generated by hifiasm (n=10 haplotypes; 30 GWAS analyses; Fig. S7,S8) and selected all genes within 500 kb of the peak SNP. Strikingly, gene content and copy number varied considerably across the assembled haplotypes (Fig. 4a). Notably, in performing GWAS to each haplotype, we also uncovered what we believe to be substantial mapping artifacts introduced by using certain references for GWAS (Supplementary Text). We found that *xmrk* is consistently present on at least one haplotype from each spotted individual and is absent in the diploid assembly generated from an unspotted male. While there was substantial copy number variation for both protein-coding genes and lnc-RNAs this region (Fig. 4a), we did not find evidence that any of these features have previously described roles in pigmentation based on a literature search. We emphasize that our sample of 1-3 long-read assemblies per phenotype makes this analysis somewhat qualitative by nature, and more individuals are required to strengthen the association between phenotype and *xmrk* presence-absence variation as well as to definitively rule out other candidates for the patterning gene (referred to as Mdl in previous literature).

**Figure 4.**
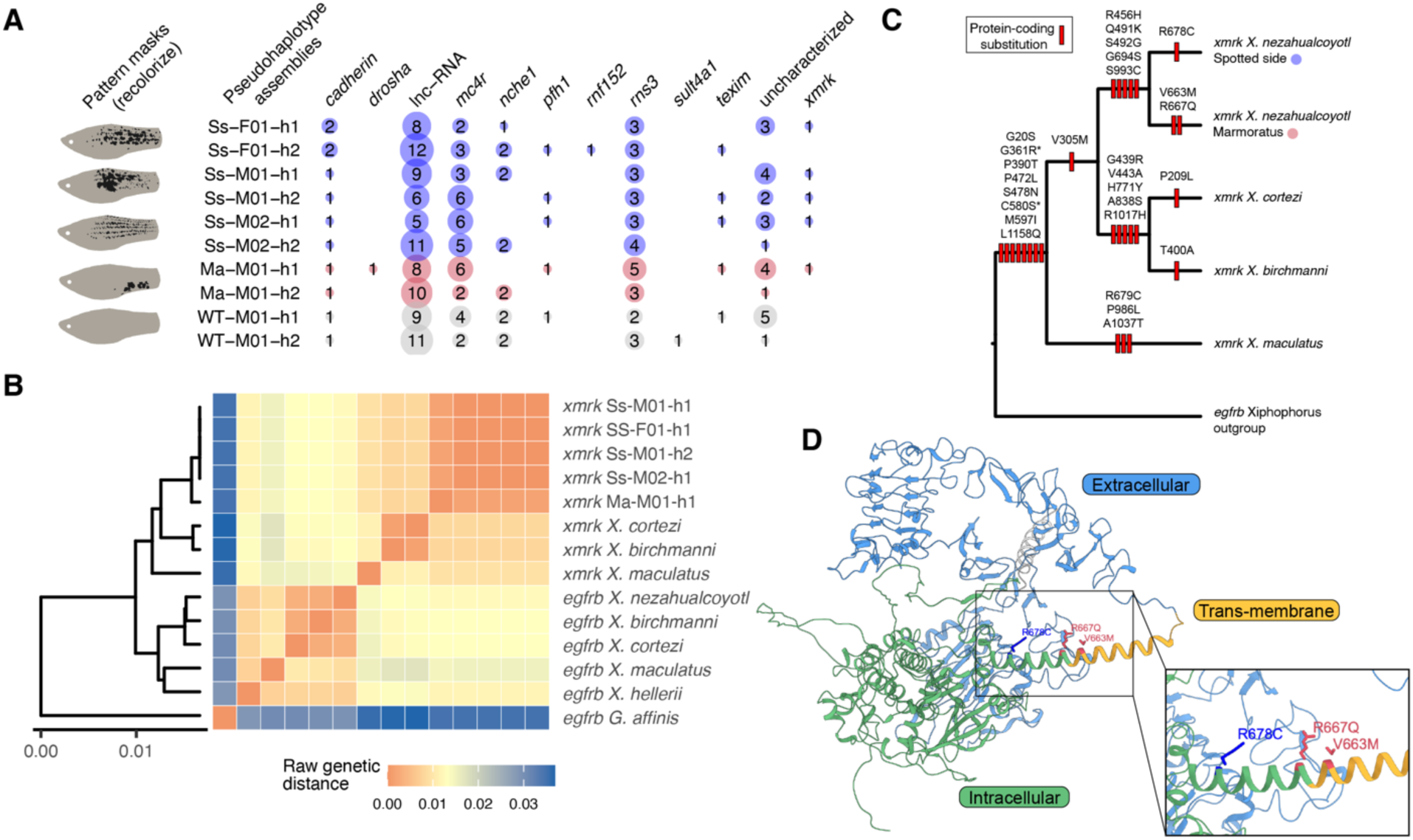
*xmrk* is associated with macromelanophore traits in *X. nezahualcoyotl* and harbors derived protein-coding changes. **(a)** Copy number variation of predicted protein-coding genes and lnc-RNAs within 500 kb of the GWAS peak for each haplotype, listed in alphabetical order due to gene order differences between haplotypes. Numbers denote gene copy number and colors denote phenotype (blue—spotted side; red—marmoratus; grey—unspotted). The lnc-RNAs and uncharacterized protein coding genes are each collapsed into a single category. *xmrk* is the only unique protein-coding feature consistently associated with spotted individuals in our sample. **(b)** Pairwise substitutions between *xmrk* and *egfrb* coding sequences indicate that the genes annotated as *xmrk* in *X. nezahualcoyotl* represent the same ancestral *xmrk* sequence as in other *Xiphophorus*. **(c)** Non-synonymous amino acid substitutions plotted on the *xmrk* tree. Stars denote amino acid substitutions that were previously identified as driving constitutive, ligand-independent activity in *xmrk*. *X. nezahualcoyotl* have accumulated five derived substitutions in *xmrk* since their split from *X. cortezi* and *X. birchmanni*. Marmoratus has two private substitutions and spotted side has one. **(d)** AlphafoldII prediction of *X. maculatus xmrk* structure, colored by domain annotations queried from UniProt. The derived substitutions associated with spotted side (blue) and marmoratus (red) are in or near the predicted trans-membrane domain of the protein.

### Synonymous and non-synonymous coding variation in xmrk

To confirm the identity of the gene we putatively classified as *xmrk* in *X. nezahualcoyotl*, we first extracted putative *xmrk* and *egfrb* sequences from the *X. nezahualcoyotl* genomes and also known sequences from other published *Xiphophorus* genomes (Table S6). We then calculated pairwise divergence between all sequences and constructed a phylogenetic tree based on UPGMA hierarchical clustering (Fig. 4b). All putative *xmrk* sequences from *X. nezahualcoyotl* clustered with known *xmrk* sequences from other *Xiphophorus* species, while the putative *egfrb* sequences from *X. nezahualcoyotl* clustered with confirmed *egfrb* sequences. Past research has shown that relative to the ancestral gene *egfrb*, *xmrk* has several identified amino acid changes to the extracellular domain [44,46,49]. To confirm this for our sequences, we translated the predicted coding sequences and constructed an *xmrk* phylogenetic tree (Fig. 4c) with each substitution placed on the corresponding branch following a simple parsimony approach. At the protein level, we found that all *X. nezahualcoyotl xmrk* sequences share the eight amino acid substitutions that distinguish all *xmrk* from *egfrb*, including the predicted substitutions important for constitutive activity (G361R and C580S) in the extracellular domain [62,63].

We next explored variation in the coding sequence and predicted proteins specific to *X. nezahualcoyotl*. We found five non-synonymous derived substitutions shared among all *X. nezahualcoyotl xmrk* sequences and absent in their sequenced relatives. Three of these shared substitutions (R456H, Q491K, and S492G) fall within conserved extracellular domains. The four *xmrk* sequences derived from spotted side individuals had identical coding sequences and differed from the single marmoratus *xmrk* sequence at six nucleotides, corresponding to three non-synonymous amino acid substitutions. Two substitutions (V663M and R667Q) are derived in marmoratus, and one (R678C) is derived in spotted side. The marmoratus-derived substitutions appeared fixed within the haplotype based on an analysis of pooled low-coverage data from the GWAS sample (Fig. S9). The three amino acid substitutions segregating in *X. nezahualcoyotl* each occur in a predicted alpha helix, part of which is within the transmembrane domain (Fig. 4D). To obtain initial insight into potential effects of these amino acid substitutions, we calculated their Grantham distances, which consider amino acid composition, polarity, and molecular volume [64]. The two derived substitutions within the marmoratus haplotype have low Grantham scores (21 and 43, respectively), suggesting a more conservative impact. In contrast, the derived spotted side substitution is predicted to be non-conservative based on its Grantham distance of 180; however, it falls near the edge of the trans-membrane domain in a less confident region of the AlphafoldII predicted protein structure (pLDDT<70).

### X and Y chromosome haplotypes display extensive structural variation near xmrk

Previous crossing experiments suggested spotted side is linked to both the X and Y chromosome [28], and we therefore aimed to assign each pseudohaplotype to a X or Y chromosome. Alignments of male chromosome 21 haplotypes 1 and 2 displayed high levels of synteny, with a notable exception on the distal end in the region co-localizing with the sex-determining region identified by GWAS (Fig. 5a). Eight of the ten assemblies were gapless in this region (Fig 5a) and contained no evidence of local misassembly based on Merqury kmer analysis (Table S5). In males, one haplotype was consistently longer (27.4-28.2 Mb versus 26.4-26.8 Mb) and contained a 900 kb inversion that co-localized with the predominant signal identified in the sex GWAS. Heterozygous status of this inversion in males and its absence in females suggest that it is a Y-linked structural variant. Self-alignments suggest that the increased length of the putative Y haplotype appears to be driven largely by stretches of local repetitive content, representing ampliconic arrays flanking the chromosomal inversion (Fig. 5b). The largest array (∼1-1.5Mb), located upstream of the proximal inversion breakpoint, contained multiple copies of *mc4r*, a gene known to be ampliconic in other *Xiphophorus* species [65,66]. We found another ampliconic array (Fig. 5b, Fig. S10), which occurred downstream of the distal inversion breakpoint. This array contained distinct multi-copy genes including *cd276* and *fzd7a* and was polymorphic in both X and Y chromosomes.

**Figure 5.**
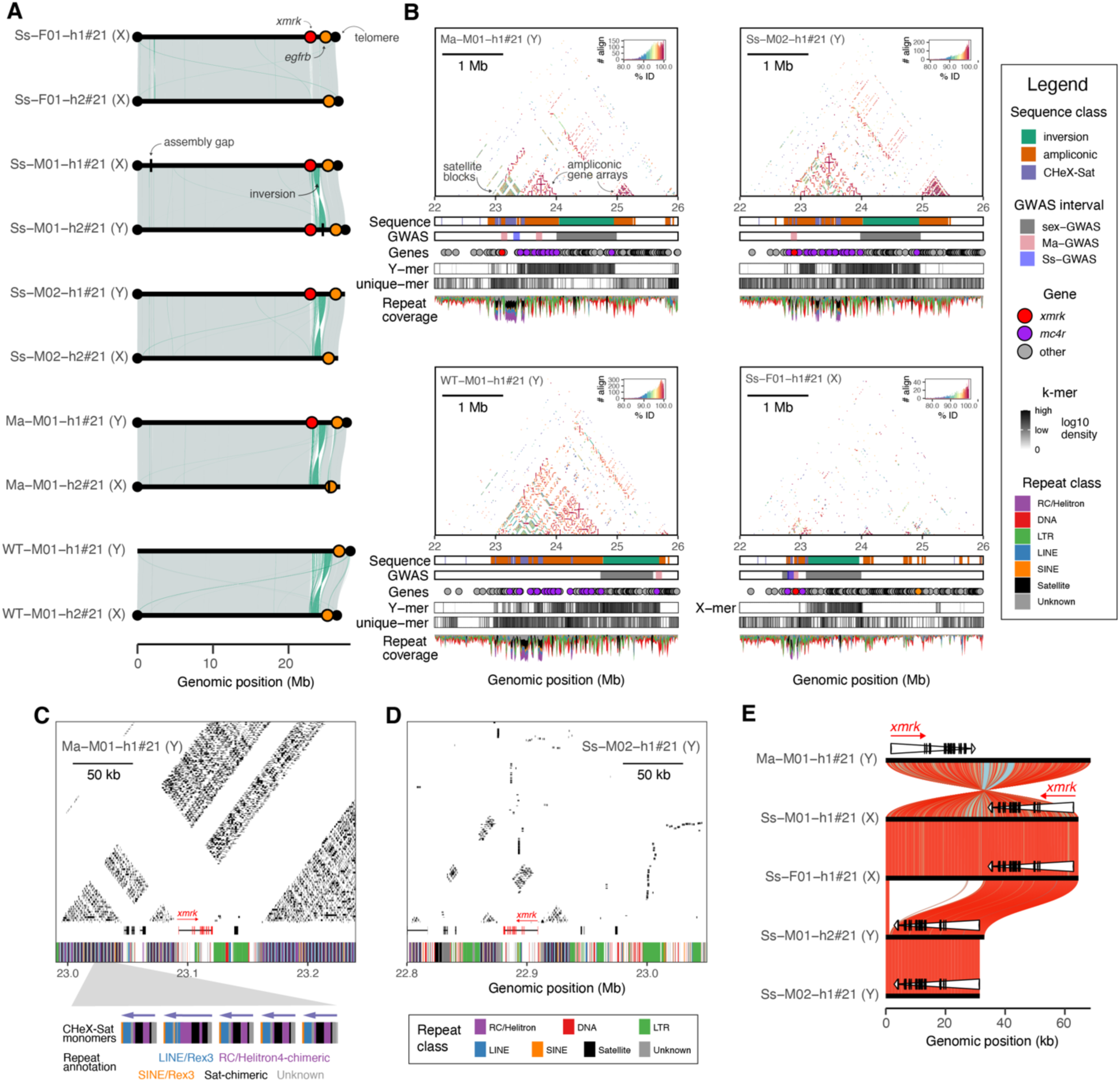
The sex-associated region is associated with extensive structural variation near *xmrk*. **(a)** Alignments between chromosome 21 haplotypes 1 and 2 are largely syntenic outside of the region associated with sex, which shows elevated rearrangement including an inversion (green) between X and Y chromosomes. Each chromosome is annotated with *xmrk* (red dots), *egfrb* (orange dots), telomeres (black dots), and gaps (black lines). Eight of the ten haplotypes are gapless across the sex-linked region. **(b)** Composite plots showing distinct structural haplotypes in 4 Mb on the sex chromosome. Top panel shows StainedGlass self-alignments, colored by percent-identity. The first two tracks show sequence class annotations and intervals defined by GWAS. 50 kb was added to each side of the GWAS interval to aid visualization. Y chromosomes are especially variable in the length of their ampliconic gene arrays and satellites arrays while these are reduced on the X chromosome. In addition to the ampliconic region containing *mc4r* (purple dots), other polymorphic ampliconic regions exist on the other side of the inverted region, which segregate in both X and Y chromosomes (see also Fig S9). Repeat content is elevated in this region. **(c)** 250kb inset showing self-alignment of “block-like” regions within ampliconic array from marmoratus Y chromosome haplotype near *xmrk*. Self-alignments show stretches with >100 bp of perfect identity. Highly self-identical regions consist of elevated repeat DNA content, which form tandemly arrayed composite satellite sequences. This 250 kb inset shows approximately half of this satellite expansion. Shown below the inset is a zoomed-in view of 5 satellite monomers, consisting of truncated Rex3 elements, a complex satellite chimeric unit, a Helitron4-satellite chimera, and several unknown elements. **(d)** 250 kb inset showing self-alignment of block like regions within ampliconic array from spotted side Y chromosome haplotype. Annotations are the same as (c). The spotted side Y chromosome has reduced satellite content in the region surrounding *xmrk*. **(e)** Alignment of *xmrk* and flanking sequence between composite satellites. *xmrk* exons and the full transcript are both represented by white triangles. *xmrk* from a marmoratus individual is inverted relative to haplotypes from spotted side individuals, and the *xmrk* haplotype from the spotted side Y chromosomes has a 30kb deletion of sequence relative to haplotypes from the X chromosomes. Alignments are colored by percent identity. Despite both being associated with Y chromosomes, the marmoratus *xmrk* haplotype is more distant from the spotted side haplotypes than spotted side on X and Y.

We next aimed to identify the Y- and X-specific regions with greater resolution than possible in the low-coverage short-read GWAS analysis. Leveraging our phased diploid assemblies, we took a complementary approach by building kmer databases of Y-specific and X-specific sequences. Mapping these sex-specific kmers back to each reference assembly localized the majority of X-mers to the inversion and Y-mers to the inversion plus the *mc4r* ampliconic array (Fig. 5b). Few X- and Y-mers fell in the region surrounding the spotted side *xmrk* haplotypes, consistent with this region recombining between X and Y chromosomes, as has been observed in other species [67]. This analysis also suggested, however, that portions of the ampliconic array containing *mc4r* are Y specific, consistent with previous work. To identify regions that had evolved more recently, we identified unique kmers present in one haplotype that were absent in all others. Interestingly, the two spotted side males had no unique kmers, and a subsequent alignment confirmed that the sex-linked region (and *xmrk* region) of the Y chromosomes were identical (Fig. S11), suggesting recent common ancestry despite distinct spotted side phenotypes. We subsequently omitted one of these Y chromosomes. We found that the regions of the Y chromosome with a high abundance of unique kmers often co-localized with structurally variable sequence, which can be visualized in the stained-glass plots (Fig. 5b). Together, this suggests presence of ongoing structural evolution in all haplotypes.

In addition to the ampliconic regions, we discovered several “block-like” patterns of self-similarity in sequences within the *mc4r* array which were sometimes adjacent to *xmrk* (Fig. 5b). An analysis of repeat content in the region showed that these blocks were highly enriched in both class I and class II repeat element families, including elements annotated as helitrons that are rare in *Xiphophorus* (∼0.06-0.07% of sequence genome-wide). A closer inspection revealed that rather than representing interspersed mobile elements, these repeats were instead truncated and tandemly repeated to form large composite satellites, which in some cases stretched several hundred kb (Fig. 5c). The most abundant satellite monomer of this type consisted of a Rex and Helitron fusion that was ∼2 kb in length. We hereafter refer to this satellite as <u>C</u>omposite <u>H</u>elitron r<u>EX Sat</u>ellite (CHeX-Sat). While CHeX-Sat sequences were generally hypermethylated, consistent with a heterochromatic state, the internal sequences annotated as Helitron-chimeric and Sat-chimeric were consistently hypomethylated (Fig. S12, Fig. S13). Prevalence of this composite satellite content varied among X (60-200 kb) and Y (200-330 kb) chromosomes, suggesting recent and likely ongoing expansions and/or contractions on both chromosomes. Nonetheless, Y chromosomes had slightly but not significantly greater CHeX-Sat content than X chromosomes in our small dataset (Wilcox test, p=0.111). The haplotypes with the most CHeX-sat content were the marmoratus and unspotted Y chromosomes although we caution these analyses are based on single haplotypes. Among the Y chromosomes analyzed, the composite satellite blocks were interspersed to varying degrees with the *mc4r* array (Fig. 5b). In the X chromosomes, where we sequenced multiple independent unspotted haplotypes, the satellite blocks formed several distinct structures ranging from nearly absent, interspersed, or occurred mainly as a single large block (Fig. S10).

Finally, we evaluated local structural differences in the region adjacent to *xmrk*. In all assemblies, *xmrk* fell upstream of the ∼900 kb inversion strongly associated with sex in the GWAS. In all spotted individuals, the *xmrk* haplotype occurred within the *mc4r* array, always between the first *mc4r* copy and all other copies (3-5 full-length copies on X chromosomes versus 10-20 full-length copies on Y chromosomes). CHeX-Sat abundance and position varied substantially near *xmrk*. At a finer scale, it appeared that *xmrk* was closely flanked by satellite blocks in all individuals (Fig. 5b). CHeX-Sat blocks near *xmrk* in the marmoratus haplotype (Ma-M01-h1#21) were substantially larger compared to spotted side (Ss-M02-h1#21) (Fig. 5c,5d). Because the satellite blocks presented formidable challenges with alignment, we extracted the region containing *xmrk* from between the nearest satellites, a region we could reliably align. We found that the marmoratus haplotype was diverged and inverted relative to all spotted side haplotypes (Fig. 5e) and contained a second nested 2 kb inversion within the first exon of *xmrk* (Fig. S14). The four spotted side haplotypes displayed high percent similarity to each other in this region (Fig. 5e). However, the spotted side Y haplotype was substantially shorter with a 30 kb region downstream of *xmrk* absent. In *xmrk* haplotypes from the marmoratus Y and spotted side X chromosomes, this region contained two LTR elements and a predicted lnc-RNA derived from the more distal element. Since these elements are polymorphic within spotted side haplotypes (between X and Y), their contribution to phenotypic variation seems less likely.

## Discussion

Despite longstanding interest in sex-linked traits, sex chromosomes have remained challenging to study, and we lack a full picture of how these dynamic regions contribute to phenotypic variation [19]. In this work, we characterize two distinct macromelanophore traits segregating in a natural population of the mountain swordtail *X. nezahualcoyotl* and investigate their genetic architecture. These patterns are allelic variants of a phenotypically diverse sex-linked locus in the genus *Xiphophorus*, whose underlying genetic architecture has remained elusive for nearly 100 years [23]. Here, we combine GWAS of hundreds of individuals with long-read sequencing of a variety of phenotypes to uncover diverse sex chromosome structure and trait architecture. Although challenges remain, including reference-bias in the short-read GWAS (see Supplementary Text), our combined approach has substantial power to resolve complex regions and identify candidate variation associated with phenotypic changes. Our work sheds light on the sex-linked macromelanophore locus and raises new insights regarding sex chromosome and trait evolution in this group. We set the stage for future studies to deploy genetic perturbations to definitively identify a macromelanophore patterning gene or variant in this region.

Advances in long-read sequencing and assembly have the potential to resolve challenging regions of genomes, which have resisted study outside of a few taxa [14,18]. Here, we generated a gapless, chromosome-level reference using only ONT ultra-long reads and hifiasm. This reference genome fully spanned all chromosomes except for two rDNA arrays, regions that were not assembled in the first human T2T genome and contain simulated sequence [68]. To our knowledge, this new reference genome represents the most contiguous and complete genome of any Poeciliid fish and underscores how far the field has come in making routine the assembly of formerly intractable regions. We analyze this gapless genome in combination with additional highly contiguous PacBio HiFi assemblies from *X. nezahualcoyotl*, in total reconstructing six X chromosomes and four Y chromosomes. Using a combination of GWAS and kmer analyses, we identify a 1-2 Mb sex-associated region with substantial divergence between—and segregating diversity within—haplotypes. The analyses revealed several features, including an inversion on the Y chromosome, ampliconic arrays containing important genes, and the presence of a newly identified class of composite satellites. These classes of sequence variants are associated with sex chromosomes in diverse taxa [9]. In particular, segmentally duplicated sequences and satellites are crucibles of genomic instability [69,70]. Little is known about composite satellites although near-T2T genomes are revealing they can be prevalent on the human acrocentric chromosomes [71]. By assembling near-complete sex chromosomes, we can gain greater understanding of variants that exist and how such changes may affect traits.

Our study combining GWAS and near-complete assemblies revealed that haplotypes containing the oncogene *xmrk* are associated with both spotting patterns. The spotted side *xmrk*-haplotype can occur on either the X or Y chromosome while we suspect that the marmoratus haplotype is Y-limited, given the absence of this phenotype in females. Most fish we sequenced with long-reads have a single *xmrk* in their diploid genome (i.e. are hemizygous), but we also sequenced a male carrying *xmrk* on both sex chromosomes. This may explain the small number of males in natural populations that have both patterns (likely a Y-linked marmoratus *xmrk*-haplotype and a X-linked a spotted side), although this remains to be tested. Notably, the position of *xmrk* within this region may impact its evolutionary dynamics. A more central position of *xmrk* nested within the satellite blocks in marmoratus versus spotted side could limit recombination between X and Y haplotypes. We speculate that reduced recombination between X and Y due to accumulation of satellite blocks may explain why the marmoratus phenotype, but not spotted side, appears to be absent in females.

In the 1990s, a debate emerged considering whether the oncogene *xmrk* controlled both melanoma and the presence of macromelanophores [44] or was merely tightly linked to Mdl and enhanced existing patterns [48]. In our analysis of the region implicated by GWAS, we did not identify any protein-coding genes or lnc-RNAs near *xmrk* with known roles in pigmentation based on a literature search. We also did not find evidence that other previously identified candidates for Mdl in other *Xiphophorus* species, including *egfrb* [36] and *myrip* [46], were associated with spotted phenotypes. However, while some pigmentation studies have famously repeatedly identified a core set of conserved candidates [72], associations with new genes are still being discovered in diverse species (e.g. *Goldentouch* in Midas cichlids [73], *Arhgap36* in orange cats [74,75]). Moreover, we cannot exclude the possibility of unannotated genes and lnc-RNAs, especially since our annotations were based on a liftover from a distinct species. Long-read RNA-sequencing data may hold special promise to identify or rule out other candidate protein-coding genes and lnc-RNAs that might be Mdl. Nonetheless, our finding that *xmrk* is present and associated with spotted phenotypes in *X. nezahualcoyotl* appears to conflict with previous reports [48]. We note, however, that the underlying model motivated by the distribution of *xmrk* and pigmentation patterns in the genus may still hold. Namely, even though we now know spotted patterns in *X. nezahualcoyotl* are linked to *xmrk*, there are still several species with macromelanophore patterns that are not thought to be associated with *xmrk*, including the dabbed patterns in *X. hellerii* [48], punctatus 1 in *X. variatus* [76], and carbomaculatus and atromaculatus in *X. birchmanni* and *X. cortezi* [28,41]. Future work studying these species may provide new insights into this longstanding debate in the literature.

Despite the uncertainties regarding if *xmrk* presence drives macromelanophore presence, our study identified coding and non-coding genomic variation associated with distinct spotted haplotypes that could play a functional role. At the protein level, there are three amino acid substitutions in *xmrk* that are predicted near the trans-membrane domain and differ between spotted side and marmoratus. Additionally, our analysis of two males with distinct spotted side phenotypes (lineated versus blotchy) revealed their Y chromosomes (and *xmrk*-haplotypes) were identical. However, the male with larger, more irregular spots had an additional, X-linked copy of a spotted side *xmrk*-containing haplotype, which could suggest a dosage-dependent relationship between the *xmrk* haplotype and pattern variation. Numerous non-coding structural differences occur near *xmrk*, including inversions, deletions, transposable element insertions, and variation in satellite content. While repetitive elements such as satellites and transposable elements have long been considered heterochromatic and deleterious, recent work has shown both TEs [77,78] and satellites can impact regulation of nearby genes [79], making their presence near the *xmrk-*haplotype (and within its potential *cis*-regulatory region) intriguing. Recent work in other Poeciliid fishes have identified helitron content associated with sex-linked selected coloration phenotypes [80], although the mechanism of action remains unknown. While outside the scope of the present study, new long-read methods profiling chromatin accessibility and DNA-protein interactions in complex repetitive regions of the genome, such as Fiber-seq [81] and DiMeLo-seq [82], may aid in identifying possible *cis*-regulatory elements in these complex sequences. Future work performing functional validation is necessary to evaluate these possibilities and distinguish association from causality in this complex genomic region.

Our work raises several new evolutionary questions related to the sex chromosomes and to sex-linked traits. It is unclear how widespread this composite satellite sequence is across fishes, and its role in the evolution in *xmrk* or a linked patterning gene (i.e. by promoting instability leading to deletion of *xmrk* or by changing the regulatory landscape near *xmrk*). Past work has pointed to elevated transposable element content as a possible driver of instability near *xmrk* [83], which theory suggests could eventually lead to sex chromosome degeneration, and it is possible that the composite satellite could play a similar role in driving sex-linked trait diversification. Moreover, the function of these two phenotypes remains unclear. Other macromelanophore traits in *Xiphophorus* may be under ecological [84] or sexual selection [85], but whether spotting phenotypes in *X. nezahualcoyotl* experience similar selective pressures is unknown. Similarly, how the two patterns are maintained within *X. nezahualcoyotl*, either by balancing selection or introgression between species (e.g. if future research finds that the genetic architecture is shared with *X. montezumae*) should be investigated. As the cost of long-read sequencing continues to fall, greater availability of new near-complete genomes in *Xiphophorus* and beyond will allow for further characterization of the evolution of sex-chromosomes, sex-linked traits, and their evolutionary history, which we anticipate will reveal even greater diversity. While the precise structure of the sex chromosomes and the genetic underpinnings of the phenotypes they encode have historically presented formidable challenges, new technologies and analytical approaches are revealing novel insights into these formerly intractable but important genomic regions.

## Materials and Methods

### Field methods

*X. nezahualcoyotl* were collected from two sites along the Río Tanchachín between 2020 and 2025 with permission from the Mexican government (Permit No. PPF/DGOPA-005/26). A GWAS sample (n=269) was collected from a single site 10 kilometers upstream of the town of Tanchachín (site code PTES; 21.913055, −99.161134). Fish were anesthetized in a buffered solution of MS-222 (100 mg/mL) and photographed on a grid background with fins spread using a Nikon d90 DSLR digital camera with a macro lens. While anesthetized, a fin clip was collected and preserved in 95% EtOH. This individual was subsequently allowed to recover and was then released. Individuals sequenced with long reads were collected from near the footbridge and from within the town of Tanchachín (site code TANC; 21.831220, −99.147773). A subset of individuals was used to establish a colony at Stanford University, where they were maintained under standard husbandry procedures (APLAC protocol #33071).

### Phenotype characterization at pattern level

To characterize the variation within and between macromelanophore spotting patterns, we performed a quantitative analysis of their spatial organization. We analyzed 50 wild-caught individuals: 31 with spotted side, 13 with marmoratus, and six with co-occurring patterns. We repeated this process for the four fish with spots that we later selected for long-read sequencing. In FIJI, we outlined one male using the polygon tool to serve as a reference (Fig. S15), and for each image in the database, we selected 12 landmarks along the fish’s outline [86,87]. We used the alignLan function from the R package patternize v0.0.5 [86] to align the images to a reference, which is used to standardize size, and exported the aligned images as PNGs. Lighting conditions were not standardized between photographs taken in the field, preventing automatic pattern quantification. We therefore manually created masks of the macromelanophore patterns on each fish using the pencil tool in Adobe Photoshop v25.11.4 [88] to mark the macromelanophore patterns with a high contrast color. We increased the exposure by 1-3x depending on lighting quality on an as-needed basis before manually creating the masks. We imported the masked images into R and generated binary masks using a modified version of the recolorize-patternize workflow [60]. We used the recolorize2 function from the R package recolorize v0.2.0 [87] with the parameters “histogram” and two bins to create color palettes of each fish and clustered all the palettes to create a “background” color and a “pattern” color. These two colors were then imposed onto each image to create binary masks. We converted the binary masks into raster objects, which contain binary matrices where each pixel is coded as 0 for the “background” color and 1 for the “pattern” color. To visualize variation between spotted side and marmoratus, we summed the raster objects and plotted heatmaps of each pattern. We excluded individuals that we classified as having both patterns from this analysis to reduce noise but included them in the principal component analysis. To further characterize spatial variation between and within the patterns, we used the patPCA function to perform principal component analysis on the pattern raster objects.

### Phenotype characterization at cellular level

We measured melanophore cell size in individuals of several phenotypic classes: fish with spotted side (n=3), marmoratus (n=3), or unspotted (n=3). Fish were anesthetized in a buffered solution of MS-222 (100 mg/mL) until cessation of movement. Initial imaging was performed using a Leica K7 Color Microscope Camera at 0.61x (300ms exposure) and 2x magnification (800ms exposure). Locally, macromelanophores inhibit presence of micromelanophores [89], and we did not find macromelanophores and micromelanophores on the same scale. We therefore also selected one scale from an unspotted region of the skin to estimate uncontracted micromelanophore size for all individuals. From fish with spotted side or marmoratus, we selected a second scale originating from a spotted location to estimate uncontracted macromelanophore size. We dissected and mounted scales on glass microscope slides (Fisherbrand ColorFrost) in PBS with two binder clip stickers to raise the coverslip height to avoid damaging the scale (1x, 7.4pH) solution. We imaged scales at 4x and 20x magnification on a BioTek Cytation 5 imaging reader in color brightfield channels. We post-processed the 20x magnification images in FIJI [90] and enhanced the contrast by 3%. On each scale, the diameter of three melanophore cells was measured. Due to the irregular shapes of the cells, we averaged three diameter measurements for each cell to estimate diameter for a given macromelanophore cell. For statistical analyses, we performed a 2-way ANOVA in R v4.5.3, setting our independent variables as melanophore type and pattern phenotype and our response variable as the mean diameter of the three melanophore cells measured per scale. In doing so, we treated the fish (not cells) as the biological replicate for all statistical analyses.

### Long-read sequencing and genome assembly

For all samples selected for long-read sequencing, fish were euthanized with an overdose of buffered MS-222 (400 mg/mL) followed by severing the spinal cord. We extracted high molecular weight DNA from flash-frozen brain tissue using the NEB Monarch HMW DNA Extraction Kit for Tissue (Catalog #T3060L, NEB, Ipswich, Massachusetts), following the manufacturer’s instructions. We quantified purified DNA on a Qubit fluorimeter (Thermo Scientific, Wilmington, DE) to measure concentration and on an Agilent 4200 Tapestation (Agilent Technologies, Santa Clara, CA) to assess fragment length distributions. We sequenced libraries with >1µg of DNA and a peak fragment length >60 kb.

To obtain a gapless, near-T2T genome for one sample—a female with spotted side—we collected ONT ultra-long data. We prepared a library using the Ultra-Long DNA Sequencing Kit V14 (SQK-ULK114), following the manufacturer’s instructions. We sequenced the library using an in-house PromethION P2 Solo equipped with a R10.4.1 flow cell. Due to lower DNA input quantity than recommendations, we prepared only a single library, and removed and reloaded this library three times after flow cell washes. We performed basecalling with Dorado (v9.1) super accuracy model with CpG modifications dna_r10.4.1_e8.2_400bps_sup@v5.0.0_5mCG_5hmCG@v3. Reads surpassing a minimum quality score filter (Q10) and length cutoff (5000 bp) were used in downstream analyses. For the four additional male assemblies, we conducted Pacific Biosciences High Fidelity Sequencing (PacBio HiFi) on a Sequel II at the Cantata Genomics Core or on a Revio at University of Washington Long-reads Sequencing Core. We also prepared an ultra-long library for one of these males, as described for the female.

We used hifiasm v0.25.0-r726 [53,91] to generate diploid assemblies for all individuals. For the female ONT assembly, we used hifiasm with the parameters --ont --dual-scaf --telo-m CCCTAA. We then scaffolded the primary assembly to a near-T2T reference from the related species *X. variatus* (Table S7) using RagTag v2.1.0 [54] with the parameters -q 50 -f 500000 --remove-small --mm2-params “-x asm20 -t 4”. As is standard for *Xiphophorus*, we assigned chromosome names and orientation based on the *X. maculatus* 5.0 reference. In the case of the near-T2T female, RagTag joined no contigs into scaffolds, but this analysis was nonetheless useful because it oriented the contigs and assigned chromosome names based on homology with *X. maculatus*. For the HiFi-only assemblies, we followed the same workflow except we omitted the --ont parameter and used RagTag to scaffold the contig-level assembly to the near-T2T *X. nezahualcoyotl* female. We assembled the genome from the final male, for which we collected both PacBio HiFi and ONT data, using hifiasm specifying the --ul parameter, which builds the assembly graph based on HiFi data and uses the ultralong data to simplify the graph. We subsequently also scaffolded this assembly to the near-T2T female.

### Generating and curating a pan-TE library for swordtail species

We used pantera v0.2.2 [58] to mine active transposable elements (TEs) from swordtail genomes. Pantera uses pangenomes to identify candidate TEs from polymorphisms between assemblies from multiple haplotypes or individuals of a species. This approach is powerful because it can identify low-copy TEs that are often missed by alternative methods, and it generates substantially less fragmented TE models compared to traditional TE mining methods [58]. This reduces the need for manual consensus refinement and redundancy reduction. To generate a TE library that could be used across *Xiphophorus*, we generated pangenomes from several published and draft assemblies from our group for several species (Table S7). To maximize sensitivity and account for possible variation in TE presence across swordtail species, we ran pantera using two different approaches. First, we used pggb v0cbe945 [92]6/8/26 12:31:00 AM with default parameters to generate independent chromosome graphs between haplotypes for each diploid genome assembly such that the number of paths in each graph was two. Then, we concatenated these graphs and ran pantera as recommended by the pantera documentation. This resulted in 1525 candidate TE models. To ensure that we also capture TEs displaying interindividual/ population variation, we generated a second batch of pggb graphs for each chromosome from all 22 swordtail haplotypes with parameter -p 80. We ran pantera again on this pangenome graph, producing another 1310 candidate TE models. We concatenated our two TE libraries and clustered them using cd-hit-est v4.8.1 [93] with flags -c 0.8 -aS .8 -G 0 -g 1 -b 500 -T 20 -M 0 -d 0 to remove redundant models. Our resulting TE library contained 1151 candidate TE models for downstream classification.

We used a combination of semi-automated and manual approaches to classify and curate swordtail TE models and generally relied on five main sources of evidence for classification. These included homology to known TEs (based on RepeatClassifier [94]) results and other homology searches using TE-AID [97] and MCHelper [95]), the presence of expected protein domains (based on scans for known functional domains from MCHelper [95] and TEtrimmer [96]), presence/absence of structural features (e.g., long terminal repeats [LTRs] and terminal inverted repeats [TIRs]), expected element termini (based on manual inspection), and target site duplication (TSD) lengths. As a first pass, we ran RepeatClassifier [94] on all candidate TE models. RepeatClassifier is a homology-based approach that prioritizes accuracy rather than sensitivity. RepeatClassifier provided classifications for 426 models but could not confidently classify the remaining 725. To classify the remaining models, we first used TE-AID [97] to annotate and visualize TE sequence structures and homology to known TEs. TE-AID produces four different types of plots which aid in TE classification. First, TE-AID retrieves all prospective copies for a given TE family by blasting the consensus sequence at the respective reference genome and plotting divergence from the consensus for all fragmented and full-length copies. Then, it plots coverage with respect to the consensus. TE-AID also produced self-versus-self dotplots for TE-consensus sequences, allowing us to assess low-complexity regions as well as hallmark features of different insertion mechanisms such as LTRs and TIRs. Finally, TE-AID produces a plot showing the locations of open reading frames (ORFs) within each TE-consensus as well as their homology to known TEs. We supplemented these homology searches with additional BLASTn and tBLASTx [59] homology searches against DFAM [98] and RepBase [99] databases.

To identify functional machinery and protein features, we ran MCHelper and TEtrimmer to annotate TE-related functional domains using available models from the PFAM and Gypsy databases [100,101]. Since specific protein machinery is associated with different known TE mechanisms, these results provided additional evidence for TE classifications. We manually examined each TE consensus sequence as well as multiple copies of each TE retrieved from blasting back to a database of swordtail genomes to identify terminal sequences and TSDs. Many TEs have hallmark termini and TSD lengths (e.g. see [99]) which enable classification. This is especially useful for nonautonomous elements that often lack functional machinery. We manually examined all TE models for multiple lines of evidence before issuing final classifications. Our resulting TE library contained 840 classified TEs.

Compared to most TEs, LTRs are relatively easy to identify systematically due to their hallmark flanking long terminal repeats. Thus, in addition to our pantera approach, which exclusively identified polymorphic TEs across swordtail genomes, we separately generated a library of all swordtail LTRs. To do this, we ran LTR_FINDER_parallel [102] on 22 concatenated swordtail genome assemblies. Then, we employed LTR_Retriever [103] to filter false positives and generate consensus sequences for Swordtail LTR elements.

### Genome annotation

After generating high quality genome assemblies *X. nezahualcoyotl*, we predicted coding and non-coding features for each scaffolded pseudohaplotype. To obtain gene models, we ran LiftOn v1.0.2 [57] using annotations from *X. maculatus*, which has been intensively studied as a cancer model [33,104]. We used BUSCO v6.1.0 [55] and the actinopterygii odb12 database to assess quality and completeness of these annotations (Table S3). *xmrk* annotations were evaluated, and we manually added a single exon missing from annotation of the marmoratus haplotypes based on the genomic sequence, which contained no substitutions relative to other *xmrk* sequences and *egfrb*. To annotate repetitive content, we ran RepeatMasker v4.2.1 [105] using a the custom, curated repeat database for *Xiphophorus* with parameters - excln -e ncbi -s -no_is -u -noisy -html -xm -a -xsmall -source. We calculated abundance of each repeat class in windows of 10 kb. We also used srf [106] to conduct an initial search for potential satellite sequences. To evaluate the occurrence of non-B DNA, we used the program nBMST [107] to identify A-phased repeats, G-quadruplexes, direct repeats, inverted repeats, mirror repeats, STRS, and Z-DNA. We identified gaps of Ns using the command seqtk gap and telomeres using seqtk telo [108]. We also identified regions with potential misassemblies using Merqury [56], based on a library of 21-mers.

To annotate various sequence classes on the Y and X chromosomes, we took several approaches. We identified inverted regions based on minimap2 alignments to the near-T2T X chromosome. Ampliconic intervals were defined based on 2 kb StainedGlass self-alignments with >97% identity, copy number >5. Ampliconic intervals were merged if within 10 kb, but only elements > 20kb were plotted for visualization purposes. CHeX-Sat sequence was defined by the diagnostic Helitron/Rex/Sat-chimeric annotations in 10 kb sliding windows. The region was defined as CHeX-sat if >25% of sequence was attributed to one of these sequences, meaning >1 monomer was present, and regions were merged if within 10 kb. To define regions in the GWAS, we selected SNPs that surpassed our significance threshold and merged SNPs if within 100 kb. Repeat content was tabulated in 10 kb sliding windows.

To identify Y and X specific regions, we generated kmer libraries from long-read assemblies using meryl [56]. We used patterns of structural variation in the diploid assemblies (presence of an inversion relative to the X chromosome from the XX female) to assign each haplotype (hap1 or hap2) as X or Y. We then created de novo 63-mer libraries of each sequence with meryl. We selected this relatively large kmer size to achieve increased specificity in highly repetitive regions, such as the sex-chromosome. Larger kmer lengths can be sensitive to sequence ends and sequencing errors, but because we generated the libraries from contiguous highly accurate assemblies (less than 1 error per 7 million bases based on Merqury QV), larger kmer sizes are suitable. We first generated a library of 63-mers that were found in all Y chromosomes (n=4) but not present in a single X chromosome (n=6), which we refer to as Y-mers. We then did the opposite, identifying kmers found in all X chromosomes and not present in a single Y chromosome (X-mers). We mapped these 63-mers back to each reference genome to identify X and Y-specific regions. Finally, for each pseudohaplotype, we generated a library of unique 63-mers, which are present in a given assembly but not found in any other assemblies, and also mapped these back to their assembly of origin. We found that the two males with spotted side were likely close relatives (Fig. S11), so we repeated this analysis after omitting the less contiguous assembly (Ss-M01-h2).

### Genome wide association studies

Because the spotting patterns are polymorphic in *X. nezahualcoyotl*, we conducted GWAS to investigate their genetic basis using 269 *X. nezahualcoyotl* individuals collected from a single site on the Río Tanchachín population in 2023 and 2025. We extracted DNA from fin-clips with the Agencourt DNAdvance magnetic bead-based purification system (Catalog #A48705, Beckman Coulter, Brea, California) following the manufacturer’s protocol but using half reactions. We prepared Tn5 tagmentation-based libraries with the Illumina DNA TDE1 Enzyme and Buffer Kit (Illumina, San Diego, CA) and subsequently purified the libraries using 18% SPRI beads. Low-coverage (1.06× average) whole-genome Illumina sequencing was conducted on an Illumina HiSeq 4000 at Admera Health Services (South Plainfield, NJ).

Following the same analytical workflow as described in previous work [26,46], we mapped all reads to the near-T2T *X. nezahualcoyotl* reference genome with bwa-mem v0.7.17 [109] and removed reads with a mapping quality score less than 30 using samtools v1.16.1 [110]. We performed the case-control GWAS with the samtools-legacy program v1.16.1 to estimate allele frequency differences between case and control individuals. For all GWAS performed, we filtered the output data to remove sites with a minor allele frequency less than 5% and retained sites with coverage between 0.5 and 2 times the mean depth of variant sites in the VCF. For spotted side, the sample population contained 75 individuals with spotted side and a control group of 181 individuals with marmoratus or unspotted phenotypes. Because marmoratus was only observed in males, we limited the marmoratus GWAS to males. This sample population contained 21 males with marmoratus and a control group of 193 spotted side or unspotted males. As the patterns have been reported to be linked to the sex chromosome [28], we also performed a GWAS for phenotypic sex (based on external morphology, i.e., presence of a gonopodium in males and gravid spot in females) using 223 males and 46 females to identify the sex-determining region.

To determine an appropriate genome-wide significance thresholds for our analyses, we used a permutation-based approach. For each of the three GWAS analyses, we randomly shuffled phenotypes between individuals, and re-ran the case-control GWAS 500 times and recorded the minimum p-value for each run. To determine our threshold for genome-wide significance, we took the lower 5% quantile of the distribution of minimum p-values. Despite differences in sample size and phenotype frequencies, we obtained similar significance thresholds for all three GWAS analyses: 2.14×10^−9^ for spotted side, 2.6×10^−9^ for marmoratus, and 2.0×10^−9^ for sex. To estimate alternate allele frequencies in each phenotypic class, we selected all SNPs that were above the significance threshold calculated for each GWAS and calculated their allele frequencies in each of the two phenotype classes in the case-control GWAS. This analysis was performed using hap 2 of the near-T2T female as the reference, which does not have *xmrk* and is thus expected to be an unspotted haplotype, to aid in interpretation of the alternate allele frequency.

To identify genes and other expressed elements that we should also consider as pigmentation candidates, we used the methods described above to perform GWAS for spotted side, marmoratus, and sex with each of the long-read assembly pseudohaplotypes as reference genomes. We used bedtools v2.30.0 [111] to select annotated protein-coding genes and lnc-RNAs within 500 kb of the lead SNP of each pattern GWAS. We selected this interval to capture potential long-range (e.g., regulatory) interactions between the significant region and other elements. In several cases, we noted substantial mismapping; namely when mapping to references where *xmrk* was absent, a second GWAS peak formed at *egfrb*. In these cases, to select a lead SNP, we omitted SNPs within 1 Mb of *egfrb* and therefore focused on the homologous region to *xmrk*. In cases where there was a GWAS signal for both spotted side and marmoratus, we selected the lead spotted side SNP. In several cases, the signal near xmrk did not exceed our simulated significance threshold, and in these cases we still selected the lead SNP even if it was not significant, for this homology analysis.

To evaluate if population structure was present in our sample and associated with either phenotype, we performed PCA analysis. Because the data were low coverage, we generated “pseudo-haploid” calls by randomly sampling a read from each variant position and assigning the allele supported by that read. Sites not covered by any reads were coded as missing. We used plink v1.90b6.5 [112] to filter sites with a minor allele frequency less than 5% or missingness greater than 75%. We then used plink to perform the PCA analysis.

### Coding variation in xmrk

To confirm the correct annotation of *xmrk* and *egfrb* in *X. nezahualcoyotl*, we queried LiftOn annotations of confirmed *xmrk* and *egfrb* sequences from published *Xiphophorus* genomes and an additional *egfrb* sequence from *Gambusia affinis* to serve as an outgroup (Table S6). We aligned the coding sequences with Mafft v7.525 [113] and used the R package Ape v5.8-1 [114] to calculate pairwise divergence between all sequences. From the resulting distance matrix, we built a UPGMA tree using the R package phangorn v2.12.1 [115]. To investigate potential functional changes in the *xmrk* protein, we generated predicted protein sequences for each coding sequence. In this alignment containing *Xiphophorus* and *Gambusia*, the *xmrk* and *egfrb* proteins are structurally conserved, except for the first exon. We again aligned the sequences with Mafft, resulting in a multi species alignment containing 14 sequences. To predict functional annotations and domains for the *xmrk* protein, we queried the UniProt database for the predicted 3D structures by alphafold2 of *xmrk* from *X. maculatus*. Plots were visualized using ChimeraX v1.12 [116] and colored by the domain annotations on UniProt. To predict the possible impacts of derived amino acid substitutions, we calculated Grantham distances using the R package grantham v0.1.4 [117].

### Genome alignment and phylogenetic analyses

We aligned whole chromosomes using minimap2 v2.28-r1209 [118] with parameters -x asm20 -c --eqx. We visualized minimap2 alignments using the plotAVA function in the R package SVbyEye [119]. We generated self-alignments using MUMMER v4.0.1 [120] with parameters mummer -maxmatch -l 100 -b -c -n -F -L and also with StainedGlass [121]. Smaller regions, including genes, coding sequences, and amino acid sequences were aligned using Mafft (v7.525). For phylogenetic analyses, we obtained predicted coding and protein sequences from NCBI, which included *Xiphophorus* species and the outgroup *Gambusia affinis* (Table S6).

As an additional confirmation of gene locations and copy number, we supplemented our annotation analyses with BLASTn [59] searches for the *xmrk* sequence and *mc4r* coding sequence. We filtered results by e-value (1×10^−10^) and percent similarity (≥93%). For *mc4r*, which is a single exon gene, we imposed an additional length filter of 700 bp to keep only full-length hits.

### Satellite analyses

We identified initial satellite candidates from the block-like signature present in the StainedGlass alignments. A further investigation of MUMMER self-alignments indicated that these regions contained stretches of high self-identity, consistent with tandem duplications. The repeat annotations of these sequences also showcased a predictable, tandemly repeated pattern of a truncated Rex3, a complex Helitron4-Rex3-immunoglobulin-satellite chimeric unit, another Helitron4-satellite chimera, and several unknown elements. Moreover, all elements were fragmented and contained no predicted proteins with homology to known TE proteins in RepeatPeps.lib. Moreover, none of these elements contained any functional domains associated with any TEs. Together, these lines of evidence suggest that these elements are tandemly repeated satellites consisting of truncated, chimeric transposable element sequences as well as additional unrelated sequences.

To perform haplotype-specific analyses of methylation, we concatenated our pseudohaplotypes and mapped native PacBio HiFi reads derived from the same individual to its matched diploid assembly using pbmm2 v1.16.0 [122] with the --CCS preset. We removed unmapped reads, supplementary, and secondary alignments from the output with the samtools flag -F 2308. We then used PacBio CpG tools v2.3.2 [123] to assign a methylation score to each CpG site. We calculated mean methylation score by each repeat class annotated within the CHeX satellite regions, many of which represent sub-elements of the composite satellite.

## Acknowledgments

We thank our colleagues Paola Fascinetto-Zago, Ben Moran, Majo Rodríguez Barrera, and Gil Rosenthal for assistance collecting fish. Stanford University provided computational support and resources for this project. We are grateful to Dorian Noel for providing illustrations. We thank Nick Altemose for pointing us to literature on human composite satellites. We are also grateful to members of the Schumer Lab, Manfred Schartl, Ben Sandkam, and Nick Altemose for critical feedback on the manuscript.

## Data Availability

All data for this study will be made fully available without restriction. Raw sequence data will be available through NCBI Sequence Read Archive (accession numbers pending). Genome assemblies will be deposited on NCBI (accession numbers pending). Phenotype data, images, and files necessary to reproduce figures in the manuscript will be made available on dryad. Scripts used for this project can be found on github (https://github.com/tododge/xnezahualcoyotl_spots/).

## Author Contributions

Conceptualization L.A.G., T.O.D.; Data curation L.A.G., T.O.D.; Formal analysis L.A.G., L.G., T.O.D.; Funding acquisition M.S.; Investigation L.A.G., L.G., J.J.B., R.S., N.B.H., K.E.H., T.R.G., G.A.P.; Methodology L.A.G., T.O.D.; Project administration D.L.P., T.O.D, M.S.; Resources M.S.; Software L.A.G., L.G., T.O.D.; Supervision K.E.H, P.H.S., D.L.P., T.O.D., M.S.; Validation L.A.G., L.G., T.O.D.; Visualization L.A.G., T.O.D.; Writing – original draft L.A.G., L.G., T.O.D.; Writing – review & editing L.A.G., L.G., R.S., N.B.H., K.E.H., G.A.P., P.H.S., D.L.P., T.O.D., M.S.

