## Supplementary Information for "Two melanic pigment patterns are associated with a sex chromosome-linked oncogene in the mountain swordtail *Xiphophorus nezahualcoyotl*"

#### Supplemental Information

#### Supplementary Text

##### Choice of reference genome and read mismapping artifacts

Long-read sequencing enables assembly and analysis of complex genomic regions [1]; however, its current high cost restricts implementation at population-level scales, particularly in non-model species. Here, we employed a mixed-sequencing approach, where long-read sequencing is performed on a subset of individuals to generate high-quality genome assemblies, and short-read sequencing is performed on the remaining individuals for reference-based analyses requiring larger samples (e.g. GWAS). However, errors can arise when mapping short-read sequences to polymorphic and complex regions [2]. One well-documented artifact is reference bias, which occurs when reads matching the reference genome map more readily to reference alleles than reads containing alternate alleles [3]. In population-scale studies, this can cause over-representation of reference alleles and under-representation of alternate alleles as variants present in the population but absent in the reference genome are missed [3–5].

A perhaps less-appreciated phenomenon that has gained some recent attention is artifacts derived from read mismapping due to true differences in sequence composition between the reference sequence and GWAS panel [1,6,7]. In principle, if portions of the genome are present in an individual selected for whole-genome resequencing, but the homologous region is absent from the reference genome, the sequence may nonetheless map to a non-homologous region if it is similar. In cases where the non-homologous region is single- or low-copy in the reference genome, reads may not be filtered out using standard approaches that rely on mapping quality scores (mapQ) or standard mappability filters. If downstream analyses rely on variant calls derived from these regions, spurious associations may occur. Read mismapping has been discussed in several contexts, including in eQTL [6] and microbiome analyses [1]. However, its contribution to GWAS with respect to trait variation in complex regions is less well understood.

To investigate the implications of reference choice when mapping short-reads to complex genomic regions, we conducted GWAS for the two spotting patterns and sex using each long-read assembly pseudohaplotype (n=10) that we generated in this study. In the main text, we report results of GWAS using the primary assembly from the near-T2T female (Fig 3) for which chromosome 21 is a T2T X chromosome that contains *xmrk*. In the supplement, we report and qualitatively compare outcomes of GWAS using references derived from the psuedohaplotypes generated from individuals with diverse phenotypes. In these instances, chromosome 21 may be an X chromosome (Fig. S7) or a Y chromosome (Fig. S8) and may or may not contain *xmrk*.

Results of GWAS for spotting patterns to alternate references vary substantially. We identified a trend that, when *xmrk* is absent in a reference genome, a significant association was often identified with its paralog *egfrb* (Fig. S7,S8), which occurs >2 Mb away on chromosome 21 but has >96% coding sequence identity. We discuss several examples to illustrate this trend.

When a reference containing *xmrk* is used (Ss-M01-h1; Fig. S7a), there was a single significant association near *xmrk* (similar to the reference in the main text; Fig 3). However, when the reference assembly lacks *xmrk*, as was true in many cases, we observed one of several possible outcomes. In some instances, the reference contains two GWAS peaks (e.g., Ss-M02-h2; Fig. S7b), one near positions flanking where *xmrk* typically. In other cases, there only a peak at *egfrb* and no evidence of other signal (e.g., Ma-M01-h2). Compared to GWAS with spotted side, marmoratus displayed a weaker signal near *egfrb*, often with only a single associated SNP. Broad trends on the X chromosomes were similar when we conducted GWAS to assemblies containing the Y chromosomes from diverse phenotypes (Fig. S8) although the peak SNPs near *xmrk* were sometimes less significantly associated.

While we do not think the inconsistency brought from using reference genomes without the gene of interest weakens the conclusion of this work that *xmrk* is associated with spotted phenotypes, these artifacts are certainly notable and a cause for concern. The case of the near-T2T female (Ss-F01), which is hemizygous for *xmrk,* is particularly illustrative. Only GWAS with the primary assembly (or hap1) containing *xmrk* identified a significant association with *xmrk*. If hap2 had been assigned as the primary assembly, and if we had not run GWAS using multiple haplotypes, it is possible this association with *xmrk* would not have been discovered. We note there has been over three decades of research on *xmrk* and a long controversy surrounding its role as Mdl, which is an unusually substantial body of work for a lineage-specific gene in a non-model species. This highlights potential challenges in identifying the drivers of associations in cases where the underlying genetic architecture is a presence-absence variant in a structurally challenging region. Recent work has proposed that the use of pangenome graphs may alleviate this issue by reducing mismapping rates [7]. However, pangenome approaches may be more limited in complex regions of the genome, such as the sex chromosomes. Thus, we urge researchers to consider how the selection of the reference genome can have large impacts on GWAS results, and suggest that further work is needed to improve tools and approaches for challenging genomic regions.

**Table S1. Long-read sequencing statistics for all samples.** We generated between 30x and 80x coverage of PacBio HiFi or Oxford Nanopore Technologies long-read data for individuals of varying phenotypes.

**Table S1. Long-read sequencing statistics for all samples.** We generated between 30x and 80x coverage of PacBio HiFi or Oxford Nanopore Technologies long-read data for individuals of varying phenotypes.

| Species | Alias | Source | Sex | Phenotype | Technology | Data (Gb) | Coverage | Read N50 | Median Read QV |
|---|---|---|---|---|---|---|---|---|---|
| *X. nezahualcoyotl* | Ss-F01 | PTES; Wild caught | F | Spotted side (Ss) | ONT UL | 57.768 | 80.79x | 57,716 | 21.762 |
| *X. nezahualcoyotl* | Ma-M01 | PTES; Wild caught | M | Marmoratus (Ma) | PacBio HiFi | 32.021 | 44.78x | 22,239 | 31.169 |
| *X. nezahualcoyotl* | WT-M01 | TANC; Lab born | M | Unspotted (WT) | PacBio HiFi | 33.171 | 46.39x | 16,283 | 32.013 |
| *X. nezahualcoyotl* | Ss-M01 | TANC; Lab born | M | Spotted side (Ss) | PacBio HiFi | 22.043 | 30.83x | 22,482 | 30.655 |
| *X. nezahualcoyotl* | Ss-M02 | TANC; Lab born | M | Spotted side (Ss) | PacBio HiFi | 34.365 | 48.06x | 21,077 | 32.306 |
| | | | | | ONT UL | 32.510 | 45.47x | 16,022 | 24.245 |

**Table S2. Long-read assembly statistics.** Contiguity (contig/scaffold N50, contig/scaffold number, gaps, % basepairs contained in longest 24 scaffolds, and number of T2T contigs), completeness (BUSCO), and correctness (Merqury QV) metrics were calculated for the primary near-T2T assembly following manual curation, as well as for both pseudohaplotypes assembled by hifiasm. Note that the mitochondrion was included in the curated near-T2T assembly. BUSCO was run on the actinopterygii odb12 database. Abbreviations: C – complete; S – complete, single copy; D – compete, duplicated; F – fragmented; M – missing; n – number.

| Assembly | Alias | Assembly size (Mb) | Contig N50 (Mb) | # contigs | Scaffold N50 | # scaffolds | Gaps | % in longest 24 scaffolds | T2T contigs | BUSCO (genome mode) | Merqury QV |
|---|---|---|---|---|---|---|---|---|---|---|---|
| Primary, curated | Ss-F01 near-T2T | 715.212 | 32.000 | 25 | 32.000 | 25 | 0 | 99.998 | 22 | C:99.4%[S:99.1%,D:0.3%],F:0.1%,M:0.5%,n:7207 | 74.3946 |
| Ss-F01 Haplotype 1 | Ss-F01-h1 | 715.062 | 31.967 | 32 | 31.967 | 30 | 2 | 99.82 | 21 | C:99.4%[S:99.0%,D:0.4%],F:0.1%,M:0.5%,n:7207 | 74.2353 |
| Ss-F01 Haplotype 2 | Ss-F01-h2 | 713.659 | 30.574 | 32 | 31.967 | 28 | 4 | 99.96 | 20 | C:99.4%[S:98.9%,D:0.5%],F:0.2%,M:0.5%,n:7207 | 78.1398 |
| Ma-M01 Haplotype 1 | Ma-M01-h1 | 713.458 | 28.966 | 81 | 31.286 | 66 | 15 | 99.16 | 6 | C:99.3%[S:98.9%,D:0.4%],F:0.1%,M:0.6%,n:7207 | 68.7963 |
| Ma-M01 Haplotype 2 | Ma-M01-h2 | 712.232 | 29.814 | 81 | 31.821 | 53 | 28 | 99.25 | 3 | C:99.3%[S:98.8%,D:0.5%],F:0.2%,M:0.5%,n:7207 | 73.278 |
| WT-M01 Haplotype 1 | WT-M01-h1 | 714.633 | 28.830 | 87 | 31.599 | 74 | 13 | 99.41 | 8 | C:99.3%[S:98.9%,D:0.4%],F:0.1%,M:0.6%,n:7207 | 70.4548 |
| WT-M01 Haplotype 2 | WT-M01-h2 | 714.861 | 28.089 | 93 | 31.842 | 66 | 27 | 99.40 | 7 | C:99.4%[S:98.9%,D:0.5%],F:0.1%,M:0.5%,n:7207 | 70.0888 |
| Ss-M01 Haplotype 1 | Ss-M01-h1 | 713.331 | 24.745 | 89 | 30.008 | 50 | 39 | 99.57 | 3 | C:99.4%[S:99.0%,D:0.3%],F:0.1%,M:0.5%,n:7207 | 69.4535 |
| Ss-M01 Haplotype 2 | Ss-M01-h2 | 714.046 | 26.824 | 74 | 30.068 | 49 | 25 | 99.40 | 4 | C:99.2%[S:98.8%,D:0.4%],F:0.2%,M:0.6%,n:7207 | 71.817 |
| Ss-M02 Haplotype 1 | Ss-M02-h1 | 715.514 | 29.801 | 54 | 32.006 | 48 | 6 | 99.56 | 12 | C:99.4%[S:98.9%,D:0.4%],F:0.2%,M:0.5%,n:7207 | 73.8304 |
| Ss-M02 Haplotype 2 | Ss-M02-h2 | 707.984 | 29.941 | 57 | 29.960 | 47 | 10 | 99.42 | 8 | C:98.7%[S:98.3%,D:0.4%],F:0.2%,M:1.2%,n:7207 | 71.9847 |

**Table S3. Assessment of LiftOn annotation quality using BUSCO**. BUSCO scores for predicted proteins based on NCBI annotation for *Xiphophorus maculatus* and for predicted proteins in *X. nezahualcoyotl* using LiftOn annotations. Most BUSCO genes annotated in *X. maculatus* have been successfully lifted over to *X. nezahualcoyotl.* BUSCO abbreviations: C – complete; S – complete, single copy; D – compete, duplicated; F – fragmented; M – missing; n – number.

| **Alias** | **Annotation method** | **BUSCO (protein mode)** |
|---|---|---|
| *GCF_002775205.1_X_maculatus-5.0-male* | Gnomon | C:99.0%[S:98.5%,D:0.5%],F:0.4%,M:0.6%,n:7207 |
| Ss-F01-h1 | LiftON | C:98.6%[S:98.1%,D:0.5%],F:0.6%,M:0.8%,n:7207 |
| Ss-F01-h2 | LiftON | C:98.7%[S:98.2%,D:0.5%],F:0.5%,M:0.8%,n:7207 |
| Ma-M01-h1 | LiftON | C:98.7%[S:98.2%,D:0.5%],F:0.5%,M:0.8%,n:7207 |
| Ma-M01-h2 | LiftON | C:98.7%[S:98.1%,D:0.6%],F:0.5%,M:0.8%,n:7207 |
| WT-M01-h1 | LiftON | C:98.6%[S:98.2%,D:0.4%],F:0.5%,M:0.9%,n:7207 |
| WT-M01-h2 | LiftON | C:98.7%[S:98.2%,D:0.5%],F:0.5%,M:0.8%,n:7207 |
| Ss-M01-h1 | LiftON | C:98.7%[S:98.3%,D:0.4%],F:0.5%,M:0.8%,n:7207 |
| Ss-M01-h2 | LiftON | C:98.6%[S:98.1%,D:0.5%],F:0.5%,M:0.9%,n:7207 |
| Ss-M02-h1 | LiftON | C:98.8%[S:98.3%,D:0.5%],F:0.4%,M:0.7%,n:7207 |
| Ss-M02-h2 | LiftON | C:98.1%[S:97.7%,D:0.4%],F:0.5%,M:1.4%,n:7207 |

**Table S4. Annotation of each assembly with pan-TE libraries.** Percent of genome annotated as repeat content (bases masked) for each long-read pseudo-haplotype using two libraries, Pantera (polymorphic elements) or Pantera plus LTR elements (polymorphic elements and fixed LTRs).

| Assembly | % masked with Pantera library | % masked with Pantera + LTR library |
|---|---|---|
| Ss-F01-h1 | 19.73 | 22.89 |
| Ss-F01-h2 | 19.79 | 22.94 |
| Ma-M01-h1 | 19.76 | 22.91 |
| Ma-M01-h2 | 19.59 | 22.75 |
| WT-M01-h1 | 19.83 | 22.97 |
| WT-M01-h2 | 19.71 | 22.86 |
| Ss-M01-h1 | 19.68 | 22.83 |
| Ss-M01-h2 | 19.82 | 23.00 |
| Ss-M02-h1 | 19.86 | 23.03 |
| Ss-M02-h2 | 19.7 | 22.83 |

**Table S5. Locations with potential errors on chromosome 21 identified by Merqury.** Kmer-containing intervals were merged by bedtools if within 1000 bp. Most errors occur within the first and last 100 kb of the chromosome. We detected no errors in the sex-linked region that was analyzed for structural variation the main text.

| Chr-21 assembly | Region start | Region stop | Error type |
|---|---|---|---|
| Ss-F01-h1#21 | 12755824 | 12755855 | Merqury |
| Ss-F01-h2#21 | 19745807 | 19745832 | Merqury |
| Ma-M01-h1#21 | 1943844 | 1943877 | Merqury |
| Ma-M01-h2#21 | NA | NA | NA |
| Ss-M01-h1#21 | NA | NA | NA |
| Ss-M01-h2#21 | 1595438 | 1595478 | Merqury |
| Ss-M01-h2#21 | 27500078 | 27500508 | Merqury |
| Ss-M02-h1#21 | 41 | 2875 | Merqury |
| Ss-M02-h1#21 | 7581 | 8252 | Merqury |
| Ss-M02-h1#21 | 9741 | 9769 | Merqury |
| Ss-M02-h1#21 | 11112 | 11433 | Merqury |
| Ss-M02-h1#21 | 47232 | 47262 | Merqury |
| Ss-M02-h1#21 | 2534467 | 2534492 | Merqury |
| Ss-M02-h1#21 | 6849462 | 6849498 | Merqury |
| Ss-M02-h1#21 | 27482710 | 27482750 | Merqury |
| Ss-M02-h2#21 | 0 | 1923 | Merqury |
| Ss-M02-h2#21 | 35031 | 35061 | Merqury |
| Ss-M02-h2#21 | 15485093 | 15485132 | Merqury |
| Ss-M02-h2#21 | 26569876 | 26569910 | Merqury |
| Ss-M02-h2#21 | 26576327 | 26576368 | Merqury |
| WT-M01-h1#21 | 28199444 | 28199480 | Merqury |
| WT-M01-h2#21 | NA | NA | NA |
| Ss-F01-h1#21 | 12755824 | 12755855 | Merqury |
| Ss-F01-h2#21 | 19745807 | 19745832 | Merqury |
| Ma-M01-h1#21 | 1943844 | 1943877 | Merqury |
| Ma-M01-h2#21 | NA | NA | NA |

**Table S6. NCBI accessions for outgroup sequences used in phylogenetic analysis.** Gambusia affinis, a relative with a high quality reference genome, was used as the outgroup for all phylogenetic analyses.

| **Species** | **Gene** | **Biotype** | **NCBI accession** |
|---|---|---|---|
| *Gambusia affinis* | *egfrb* | mRNA | XM_044104981.1 |
| *Gambusia affinis* | *egfrb* | protein | XP_043960916.1 |

**Table S7. Genome assemblies used to construct *Xiphophorus* Pantera and LTR repeat databases.** To broaden use of this repeat database and capture elements that may be fixed in *X. nezahualcoyotl*, we constructed repeat databases using available Xiphophorus genomes.

| Species | Assembly | Sex | Site | Technology | Publication | Status |
|---|---|---|---|---|---|---|
| *X. birchmanni* | xbir-COAC-S238-19-IX-23-M01 | M | Rio Coacuilco | PacBio HiFi | Dodge et al. 2024 | finalized |
| *X. continens* | xcon-NROF-20-III-25-M01 | M | Nacimiento de Rio Ojo Frio | PacBio HiFi | unpublished | draft |
| *X. cortezi* | xcor-PTHC-08-XII-21-M | M | Rio Huicihuyan | PacBio HiFi | Langdon et al. 2024 | finalized |
| *X. evelynae* | xeve-JUCH-S178-IX-23-M01 | M | Juntas Chicas | PacBio HiFi | Haghani et al. 2025 | finalized |
| *X. malinche* | xmal-CHIC-XI-20-M | M | Chicayotla | PacBio HiFi | Dodge et al. 2024 | finalized |
| *X. montezumae* | xmoz-ROJF-17-III-25-M02 | M | Rio Ojo Frio | PacBio HiFi | unpublished | draft |
| *X. multilineatus* | xmul-TABQ-S253-5-XI-24-M01 | M | Rio Tambaque | PacBio HiFi | unpublished | draft |
| *X. nezahualcoyotl* | xnez-TANC-19-I-22-M | M | Rio Tanchachin | PacBio HiFi | This work | finalized |
| *X. nigrensis* | xnig-C-NARC-18-III-25-M01 | M | Nacimiento de Rio Choy | PacBio HiFi | unpublished | draft |
| *X. pygmaeus* | xpyg-PTHC-S063-7-VIII-23-M01 | M | Rio Huicihuyan | PacBio HiFi | unpublished | draft |
| *X. variatus* | xvar-COAC-S280-21-XI-24-M01 | M | Rio Coacuilco | PacBio HiFi + ONT | unpublished | draft |

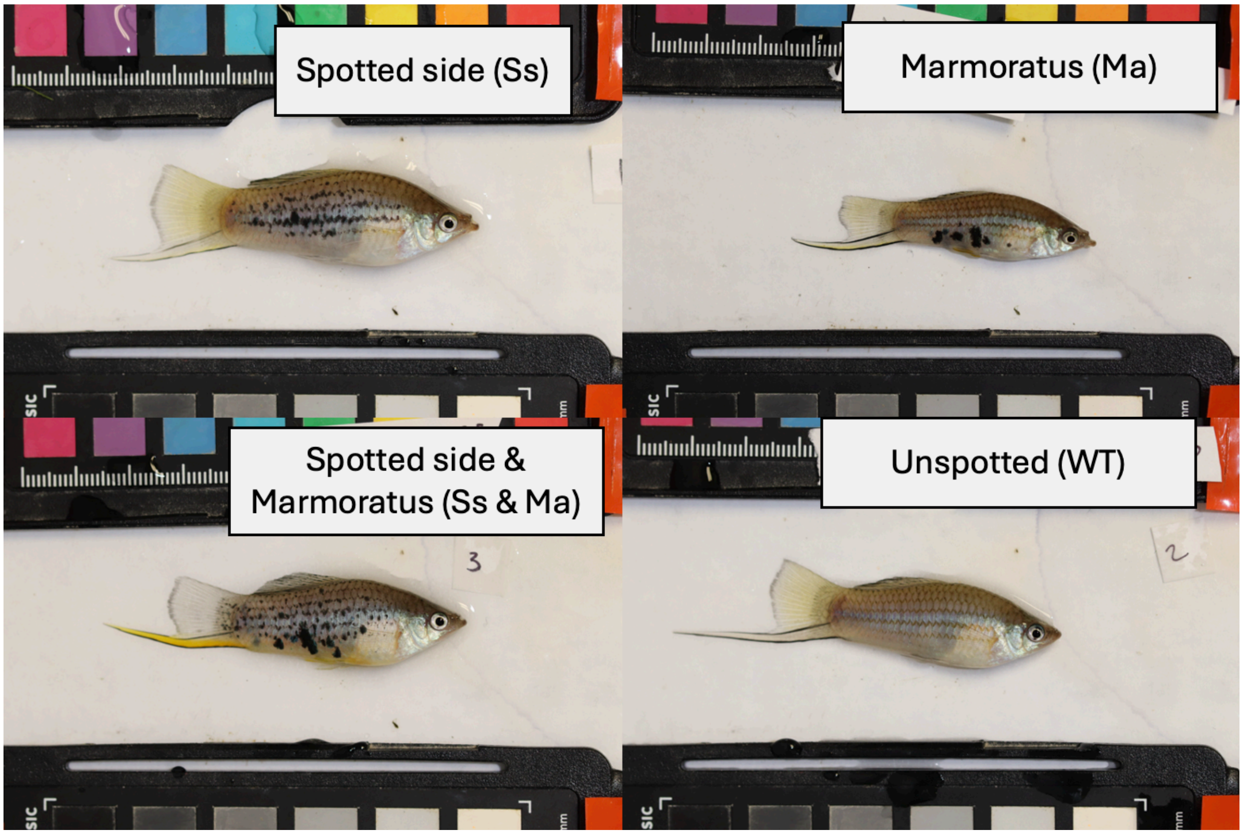


**Figure S1. Examples of *X. nezahualcoyotl* males with different spotting patterns.** Top left has spotted side (Ss), top right has marmoratus (Ma), bottom left is a male with both spotting patterns (Ss and Ma), and bottom right is unspotted (WT).

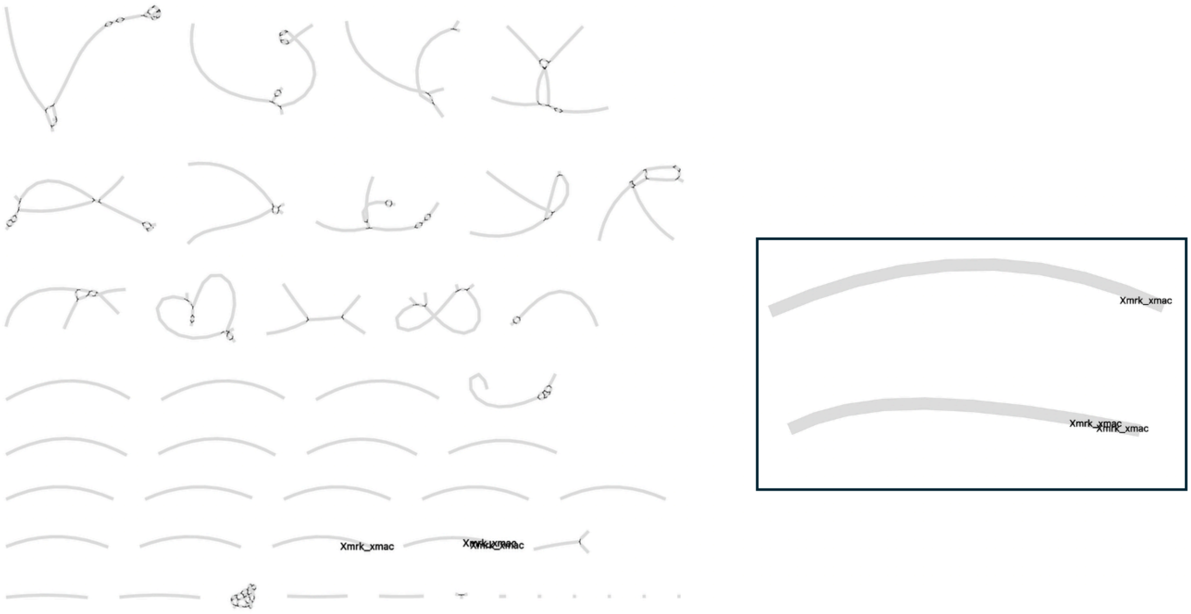


**Figure S2. Phased diploid assembly graph using ONT ultralong data**. Hifiasm unitig assembly graph for near-T2T assembly (left) and for chromosome 21 (right, black box). Blast results of *xmrk* from *X. maculatus* are shown, with one unitig containing a single hit (*egfrb*) and one unitig containing two blast hits (*egfrb* and a putative *xmrk*). Several chromosomes, including chromosome 21 (the chromosome containing the blast hits), are completely phased in the processed hifiasm unitig graph.

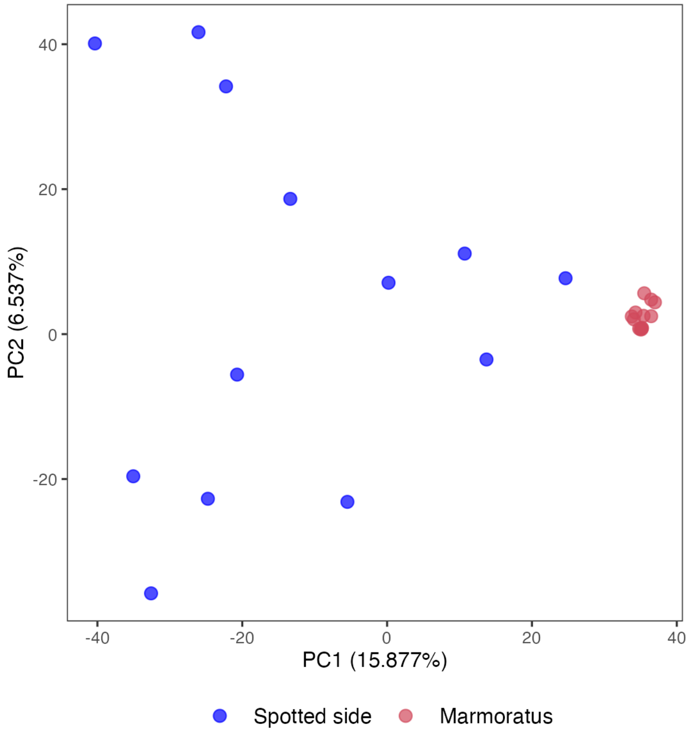


**Figure S3. Downsampled Phenotype PCA.** The pattern dataset was downsampled to include all individuals with marmoratus (n=13), and an equal number of individuals with spotted side (n=13). The separation between phenotypic clusters, and separation of the spotted side phenotype relative to marmoratus along PC2 persists after downsampling.

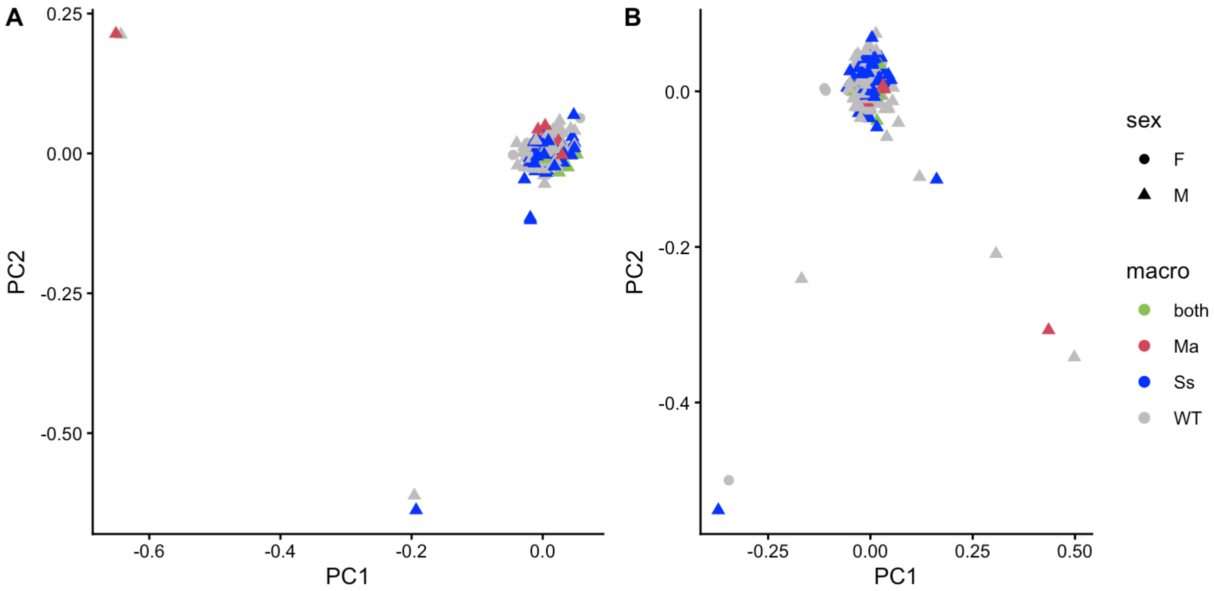


**Figure S4. Genomic PCA from the population of *X. nezahualcoyotl* used for GWAS.** We found no evidence that population structure was correlated with phenotype in this *X. nezahualcoyotl* population. 6 individuals, which represented individuals across all macromelanophore phenotype classes, appeared to be outliers. We repeated the PCA after excluding these individuals and still did not detect population structure correlated with phenotype. Note that there are substantially more males in the GWAS dataset than there are females. Ss – Spotted side; Ma – Marmoratus; WT – Unspotted.

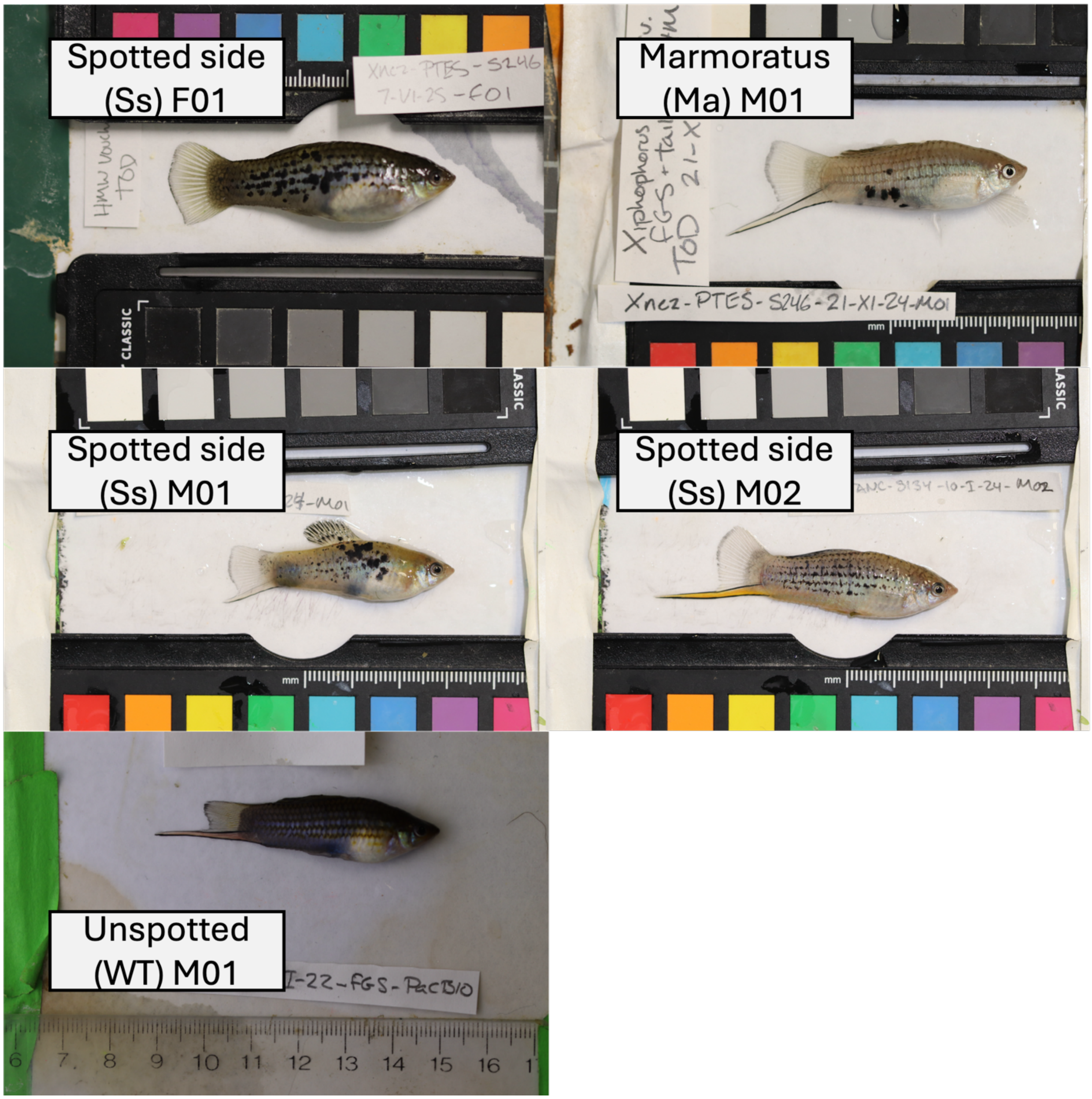


**Figure S5. Photos of the five *X. nezahualcoyotl* individuals used to produce long-read assemblies.** Top left is the near-T2T female with spotted side (Ss-F01), top right is the male with marmoratus (Ma-M01), middle are the spotted side males (Ss-M01 and Ss-M02), and bottom left is the unspotted male (WT-M01). Note that lighting conditions are not standardized between photographs, and WT-M01 was photographed before flash was implemented as part of the standard operating procedure.

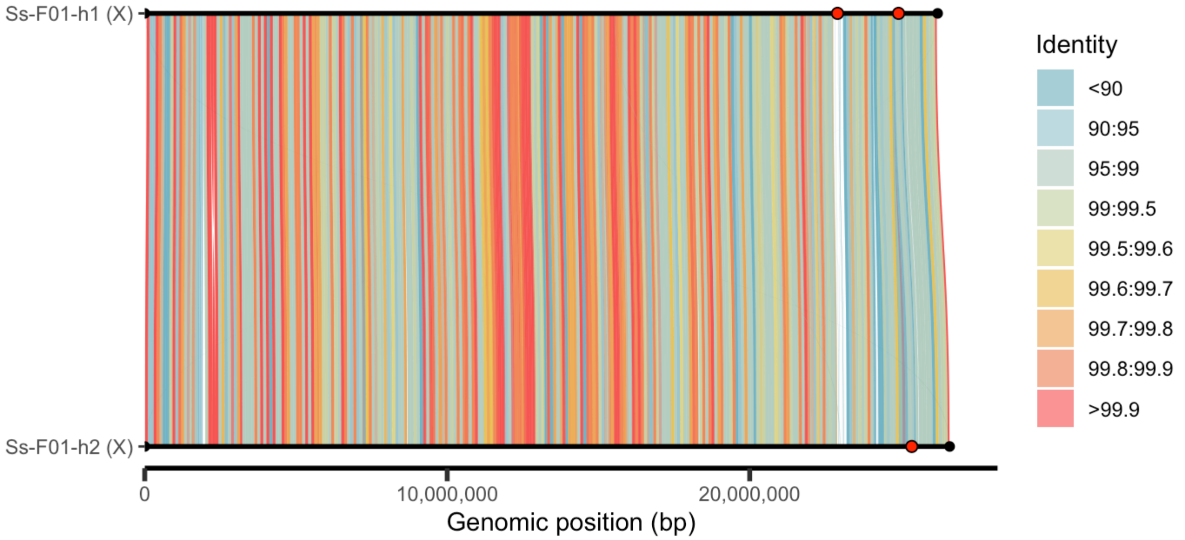


**Figure S6. SVbyEye alignments of gapless *X. nezahualcoyotl* X chromosomes.** Red dots represent *xmrk*/*egfrb* blast hits and black dots represent telomeres. The region containing *xmrk* does not align between haplotypes (white). Alignments are colored by percent identity in 100 kb bins. Note that these percent identity calculations treat gaps in the alignment as mismatched basepairs.

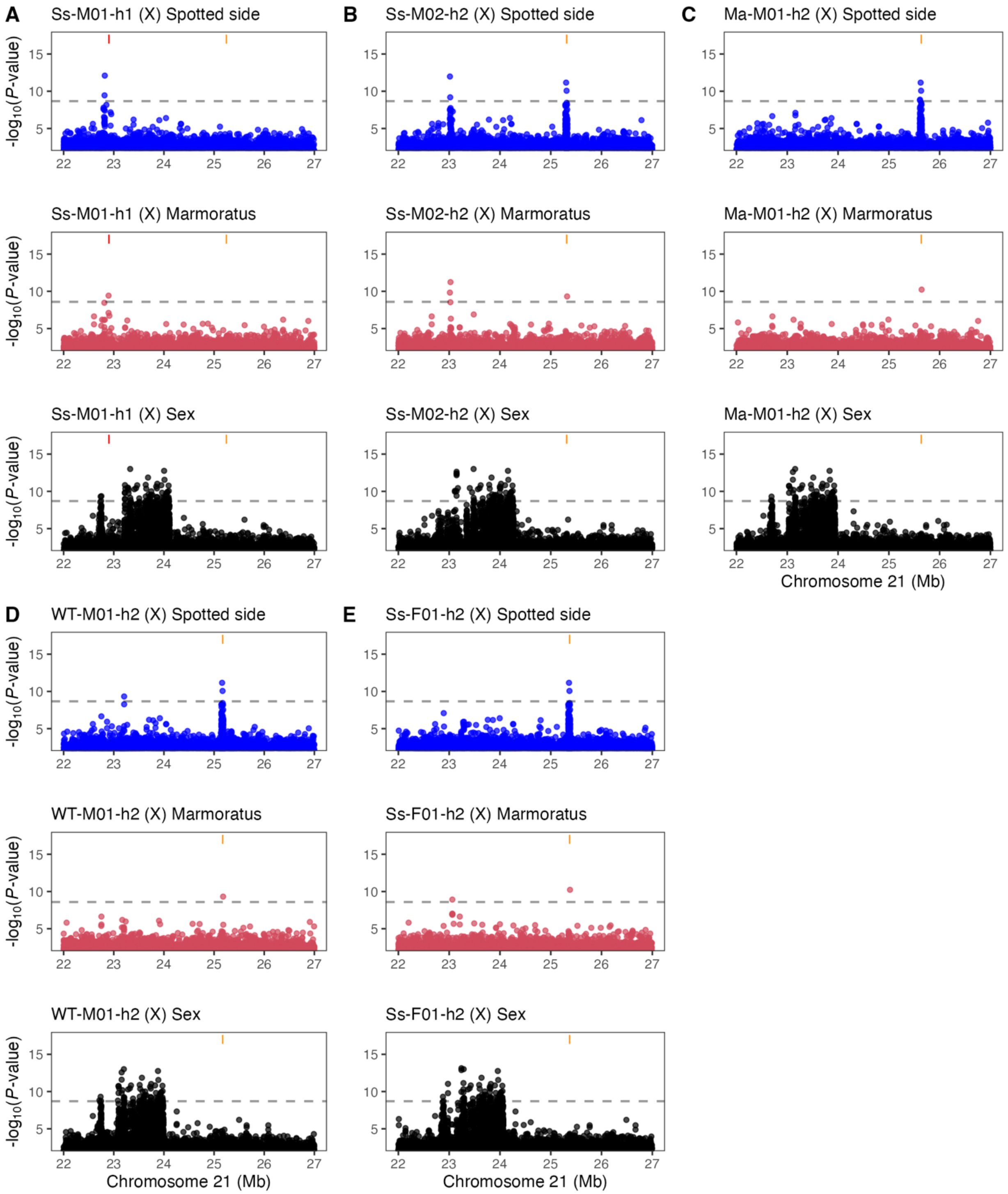


**Figure S7. GWAS results from analyses mapping data to different reference genomes, focusing on the pseudohaplotype that contains the X chromosome**. All plots span from position 22 to 27 Mb on chromosome 21, and the y-axis shows -$\log_{10}$(p-value). Blue Manhattan plots are GWAS results for spotted side, pink are for marmoratus, and black are for sex. The reference haplotype **(a)** Ss-M01-h1 contains both *xmrk* (red) and *egfrb* (orange), and the remaining haplotypes **(b)** Ss-M02-h2, **(c)** Ma-M01-h2, **(d)** WT-M01-h2, and **(e)** Ss-F01-h2 only contain *egfrb* (orange).

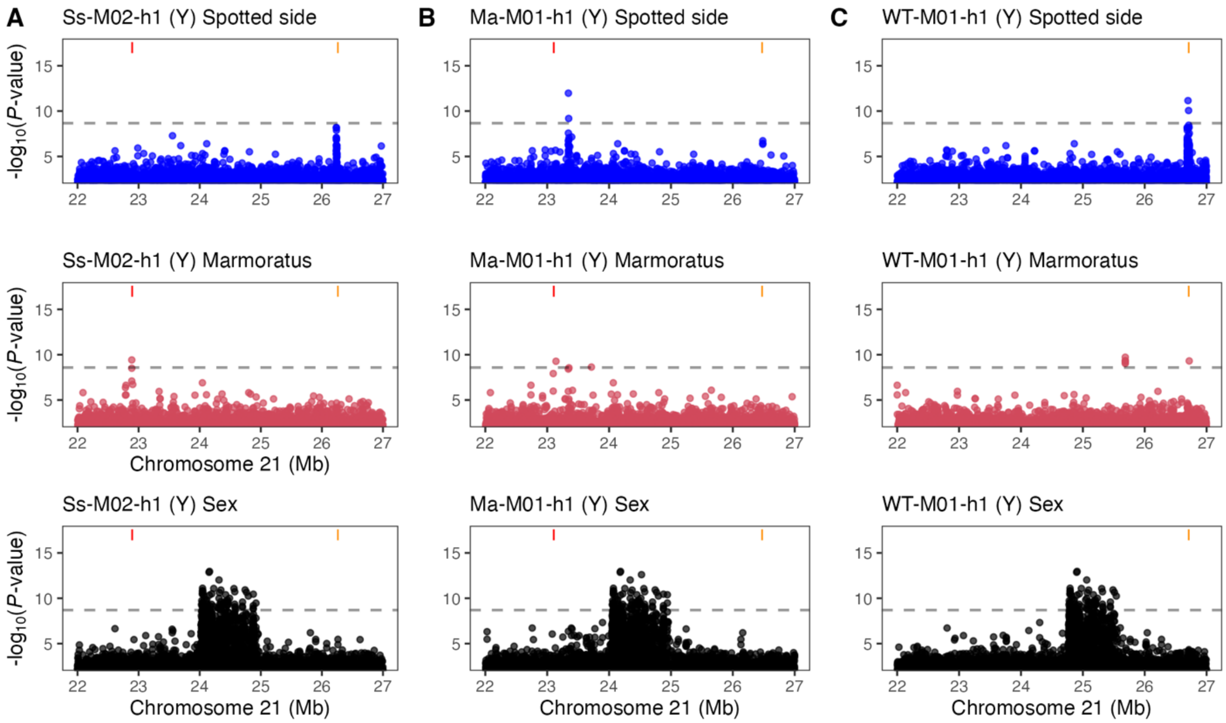


**Figure S8. GWAS results from analyses mapping data to different reference genomes, using the pseudohaplotype that contains the Y chromosome**. All plots span from position 22 to 27 Mb on chromosome 21, and the y-axis shows -$\log_{10}$(p-value). Blue Manhattan plots are GWAS results for spotted side, pink are for marmoratus, and black are for sex. The reference haplotypes Ss-M02-h1 **(a)** and Ma-M01-h1 **(b)** contain both *xmrk* (red) and *egfrb* (orange), and the reference haplotype WT-M01-h1 **(c)** only contains *egfrb* (orange).

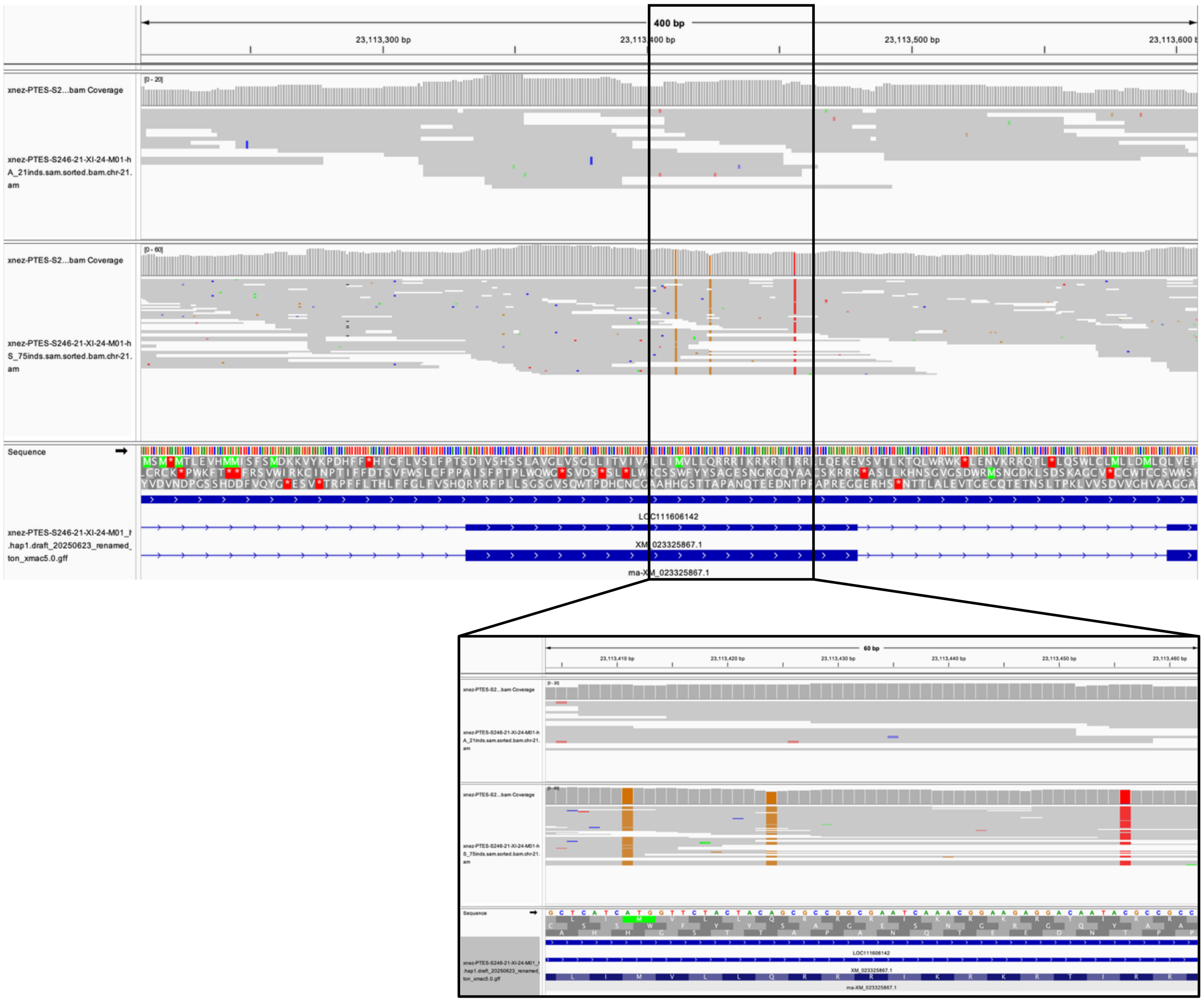


**Figure S9. Derived SNPs in marmoratus *xmrk* appear fixed in analysis of pooled sequencing data from marmoratus individuals.** Integrative Genomics Viewer showing *xmrk* sequences from pooled whole-genome short-read data mapped to the marmoratus Y chromosome. The three non-synonymous SNPs identified in the marmoratus and spotted side long-read assemblies appear to be fixed in each haplotype at the population-level.

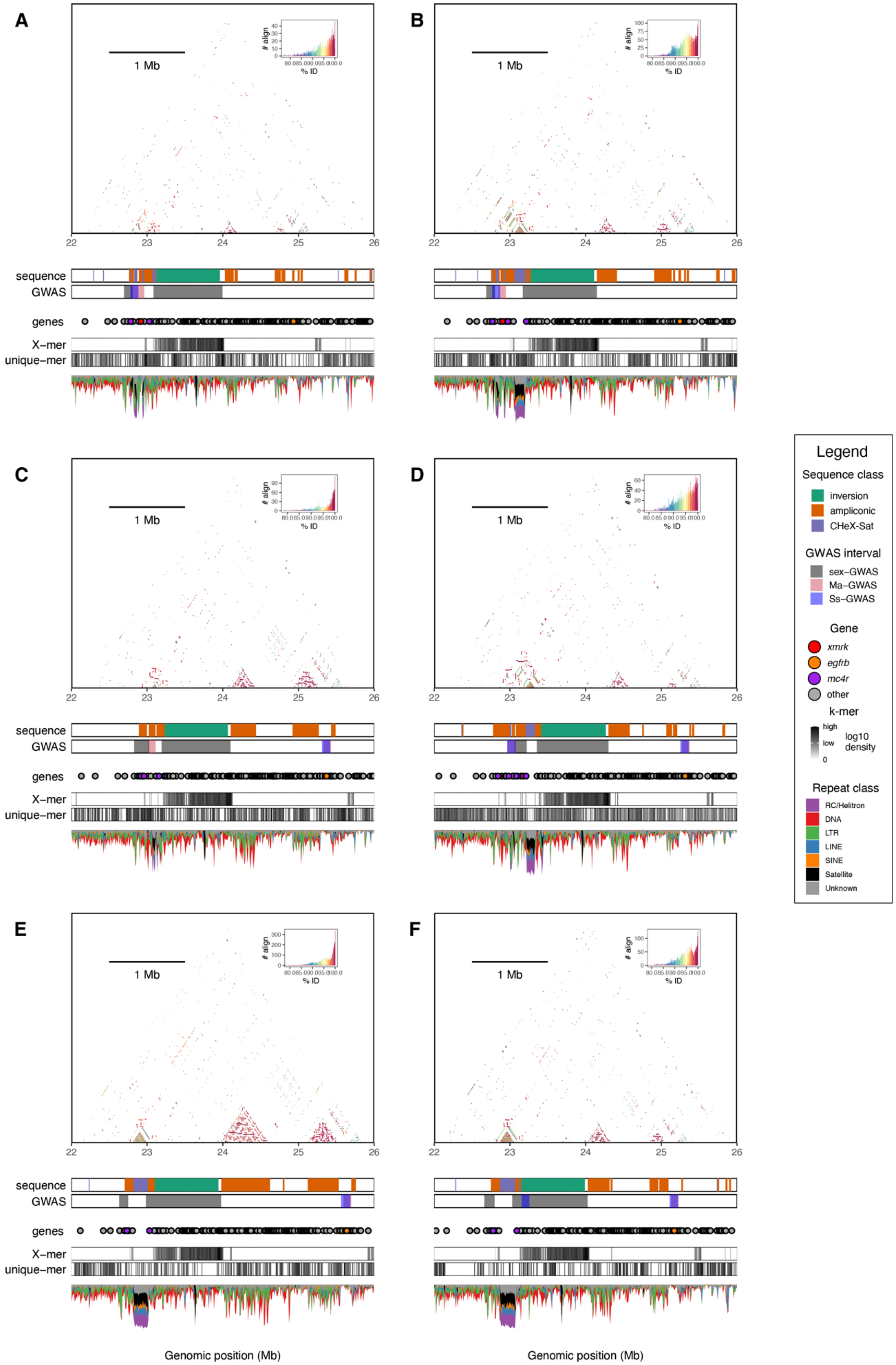
A
B
C
D
E
F
1 Mb
### align
% ID
80.0 85.0 90.0 95.0 100.0
22
23
24
25
26
sequence
GWAS
genes
X−mer
unique−mer
Genomic position (Mb)
Legend
Sequence class
inversion
ampliconic
CHeX-Sat
GWAS interval
sex−GWAS
Ma−GWAS
Ss−GWAS
Gene
xmrk
egfrb
mc4r
other
k-mer
high
low
0
log10 density
Repeat class
RC/Helitron
DNA
LTR
LINE
SINE
Satellite
Unknown

**Figure S10. Combined structural variant plots of all X chromosomes. (A)** SspF01-h1 (*xmrk*), **(B)** Ss-M01-h1 (*xmrk*), **(C)** Ss-F01-h2 (non-*xmrk*), **(D)** Ss-M02-h2 (non-*xmrk*), **(E)** Ma-M01-h2 (non-*xmrk*), **(F)** WT-M01-h2 (non-*xmrk*). For the X chromosomes in this species, repeat content (ampliconic genes and satellites) and *xmrk* presence do not appear to be clearly linked. Several X chromosomes have non-mc4r ampliconic gene expansion on the other side of the inversion, with Ma-M01-h2 having the largest. Size and number of satellite blocks also vary among X chromosomes, with Ss-F01-h1 and Ss-F01-h2 having small satellite blocks, Ss-M01-h1 and Ss-M02-h2 having a more interspersed structure, and Ma-M01-h2 and WT-M01-h2 both having one very large block.

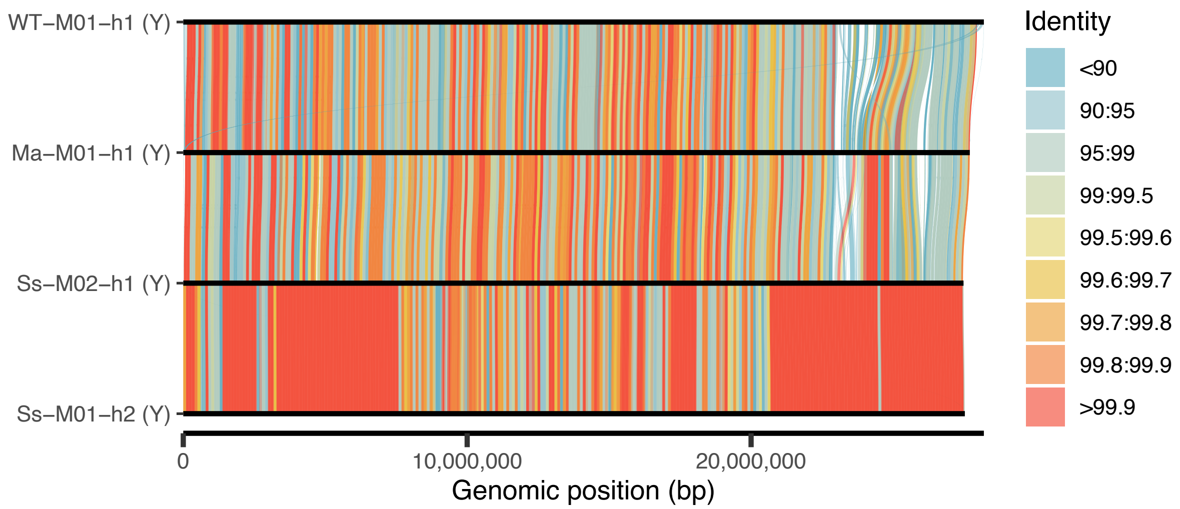


**Figure S11**. **The spotted side Y chromosomes are likely derived from close relatives.** SVbyEye alignments of *X. nezahualcoyotl* Y chromosomes colored by percent identity in 100kb bins. Note that these percent identity calculations also treat gaps in the alignment as non-matching basepairs. Ss-M02-h1 (Y) and Ss-M01-h2 (Y) were sourced from lab born males in the same tank. They have several megabase-long stretches of 100% identity, including at the end of the sex chromosome covering the sex determining region. This stretch of perfect alignment is interrupted only by a scaffold gap present in the Ss-M01-h2 assembly. These long stretches of perfect identity suggest that these two males are close (potentially first-degree) relatives, that share the sex-determining portion of the Y chromosome due to identity by descent, and therefore these two Y chromosomes should not be considered independent.

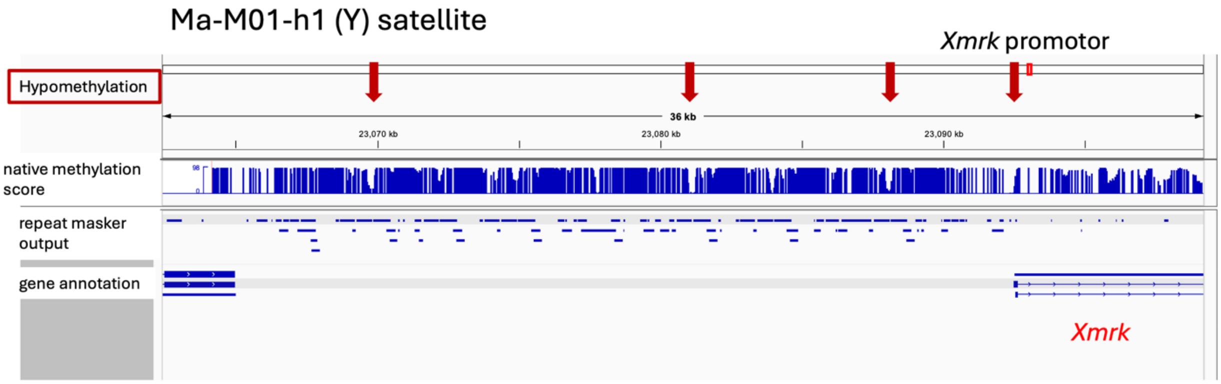


**Figure S12. Methylation status within CHeX-sat sequences adjacent to *xmrk***. PB-CpG-tools methylation scores derived from native long-read data from adult male brain tissue of the marmoratus male. The composite satellite region is generally hypermethylated except for at the boundary of sub-elements annotated as Sat-chimeric and RC/Helitron4-chimeric. The *xmrk* promotor is also hypomethylated.

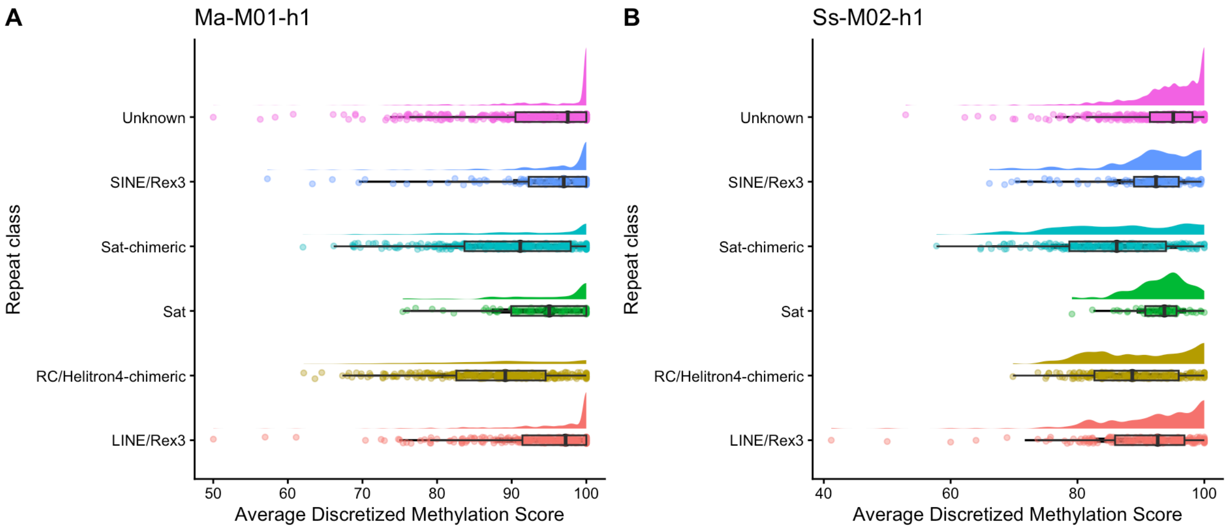


**Figure S13. Methylation status within CHeX-sat sub-elements on the sex chromosome**. PB-CpG-tools methylation scores derived from the CHeX-sat sequences from the Y chromosome of one marmoratus and one spotted side individual. Elements of the composite satellite show reduced methylation in sub-elements annotated as Sat-chimeric and RC/Helitron4-chimeric.

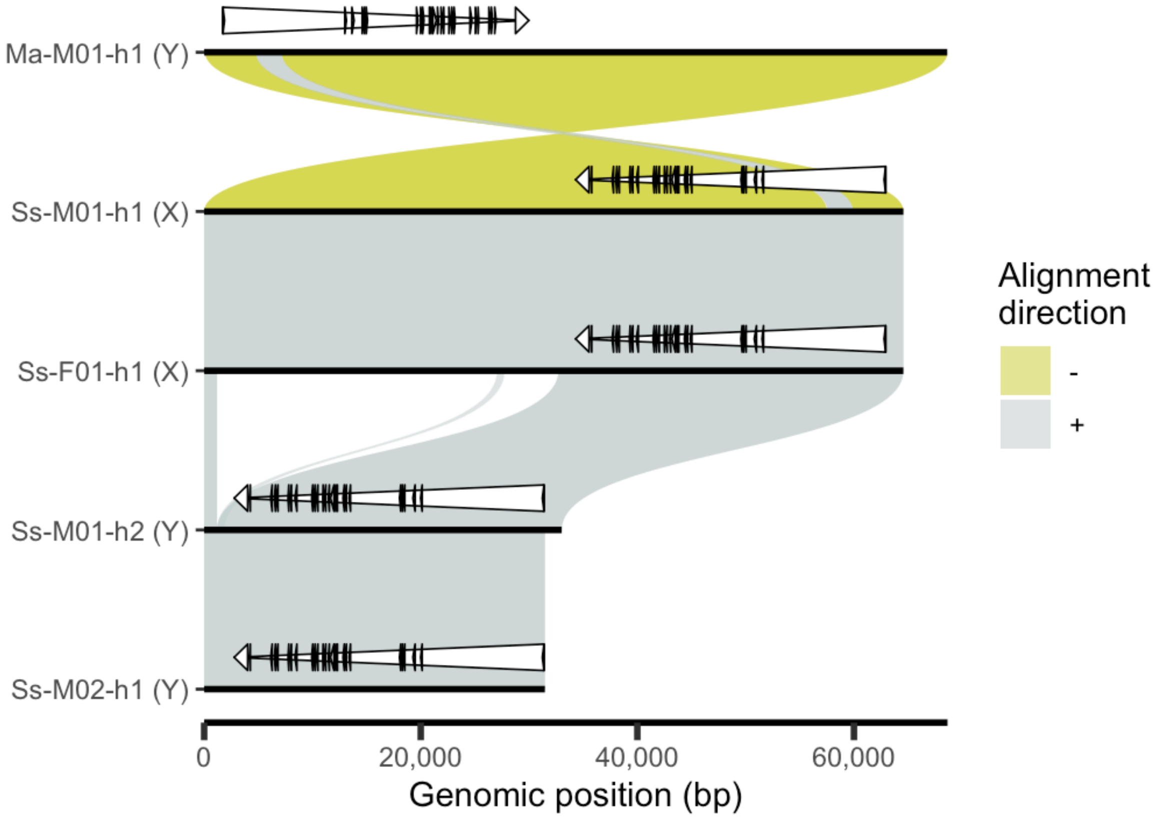


**Figure S14**. **SVbyEye alignments *X. nezahualcoyotl xmrk* sequencies colored by orientation.** The marmoratus haplotype is inverted relative to all other haplotypes and contains a nested 3 kb inversion within the first intron of *xmrk*. *xmrk* exons and the full transcript are both represented by white triangles.

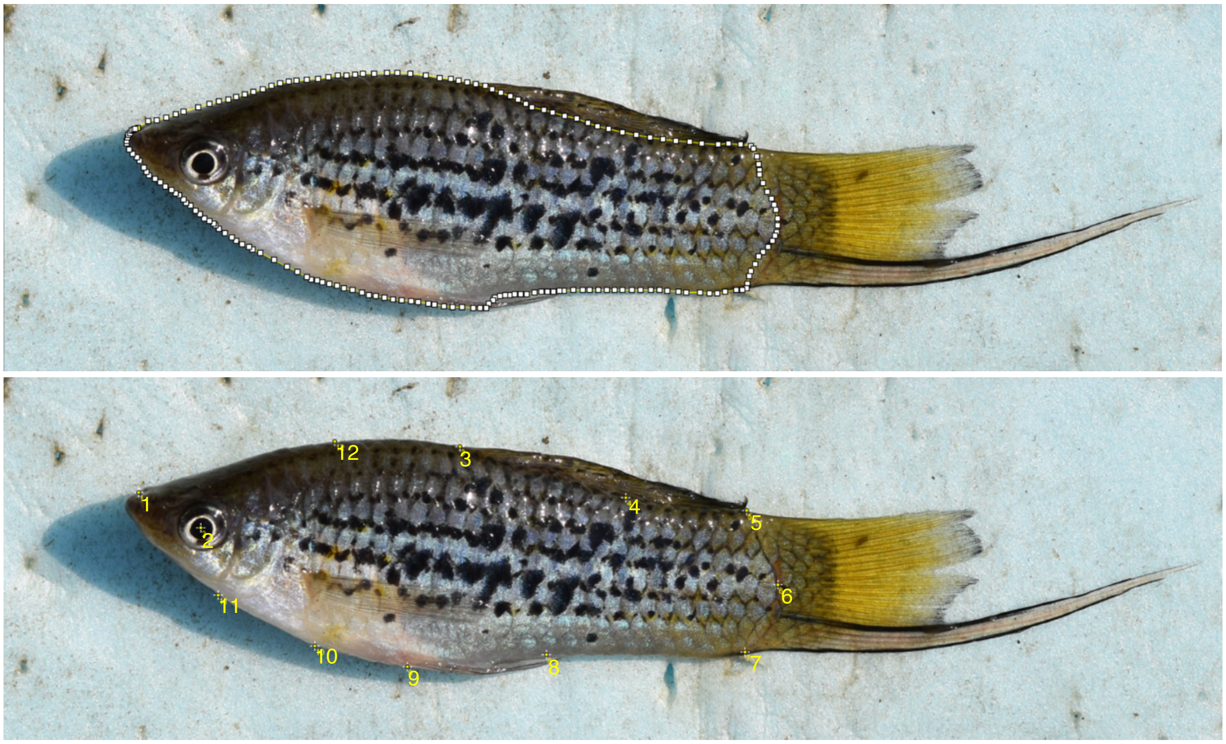


**Figure S15. Reference images for recolorize/patternize workflow**. One male was selected as a “reference” to align all images. Top shows the reference outline made using the polygon tool in FIJI. Bottom shows the 12 landmarks placed on every individual for alignment.
